# ID1 functions as an autonomous phosphate (Pi) regulator upstream of the miR399-*ZmPHO2* signaling module in maize

**DOI:** 10.1101/2022.07.29.502052

**Authors:** Xufeng Wang, Dan Yuan, Yanchun Liu, Yameng Liang, Juan He, Xiaoyu Yang, Runlai Hang, Hong Jia, Beixin Mo, Feng Tian, Xuemei Chen, Lin Liu

## Abstract

The macronutrient phosphorus is essential for plant growth and development. Plants have evolved multiple strategies to increase the efficiency of Pi acquisition to protect themselves from Pi starvation. However, the crosstalk between Pi homeostasis and plant development remains to be explored. Here, we report that overexpressing microRNA399 in maize induced a premature aging syndrome after pollination. Knockout of *ZmPHO2*, one of the miR399 targets, resulted in a similar premature aging phenotype. Strikingly, we found that INDETERMINATE1 (ID1), a floral transition regulator, inhibits the transcription of *ZmMIR399* genes by directly binding to their promoters, alleviating the repression of *ZmPHO2* by miR399 and ultimately contributing to the maintenance of Pi homeostasis in maize. Unlike *ZmMIR399* genes, whose expression is induced by Pi deficiency, *ID1* expression is independent of the Pi status, indicating that *ID1* is an autonomous regulator of Pi homeostasis. Furthermore, we found that *ZmPHO2* was under selection during maize domestication, resulting in a more sensitive response to phosphate starvation in temperate maize than in tropical maize. Our study reveals a direct functional link between Pi-deprivation sensing by the miR399-*ZmPHO2* regulatory module and plant developmental regulation by ID1.

## Introduction

Phosphorus is present in every living cell and its functions cannot be substituted by other nutrients (Marschner, 1995). As one of 17 essential nutrients, phosphorus is required for almost all developmental processes in plants and is considered one of three primary macronutrients, along with nitrogen and potassium. Nevertheless, most cultivated land worldwide is poor in phosphorus availability, and levels of inorganic orthophosphate (Pi), which plants take up through their roots, are suboptimal in soils for vegetative growth and crop productivity (López-Arredondo et al., 2014, Kirkby & Johnston, 2008, Raghothama, 1999). Along with other fertilizers, Pi must be routinely applied to farmland to support healthy crop growth. However, phosphate fertilizers used for agriculture are mostly produced from mined rock phosphate, which is nonrenewable and predicted to decline in availability before the end of this century (Cordell et al., 2011, Vance et al., 2003, Baker et al., 2015). Thus, adapting crop plants to soils with a restricted supply of Pi is vital if high crop yields, which are necessary to satisfy the food requirements of a continuously increasing world population, are to be maintained.

More Pi-efficient plants could reduce the requirement for Pi fertilizers, thereby ameliorating their overuse while concurrently enhancing yield. The investigation and manipulation of genes involved in phosphate acquisition, remobilization and metabolism is a potential strategy for the development of Pi-efficient crops, and might facilitate agricultural sustainability. To date, a series of Pi starvation-related genes and regulators have been characterized in the model plant Arabidopsis (*Arabidopsis thaliana*), including transcription factor genes such as *PHOSPHATE STARVATION RESPONSE1* (*PHR1*) (Rubio et al., 2001, Maeda et al., 2018), *PHR1-LIKE1* (Bustos et al., 2010), *WRKY75* and *ZAT6* (Devaiah & Raghothama, 2007, Devaiah et al., 2007a, Devaiah et al., 2007b), the long non-coding RNA *IPS1/At4* (Franco-Zorrilla et al., 2007, Shin et al., 2006), microRNA399 (miR399) (Chiou et al., 2006, Bari et al., 2006, Pant et al., 2008) and members of the *PHO* gene family (Stefanovic et al., 2007, Aung et al., 2006). In particular, *PHO2* (*PHOSPHATE2*), which encodes a ubiquitin-conjugating E2 enzyme (UBC), is known to be a miR399 target and is involved in Pi homeostasis (Bari et al., 2006). Under Pi-sufficient conditions, *PHO2* targets phosphate transporters for poly-ubiquitination and degradation, thereby maintaining optimal Pi uptake (Sega et al., 2020, Liu et al., 2012, Li et al., 2017). In contrast, Pi-deficiency stress induces miR399 accumulation, which downregulates the expression of *PHO2* (Bari et al., 2006) by transcript cleavage, in turn increasing the expression of phosphate transporters for the enhancement of Pi acquisition and translocation (Aung et al., 2006). The miR399-*PHO2* module also regulates Pi homeostasis in rice (*Oryza sativa*), maize (*Zea mays*), wheat (*Triticum aestivum*) and other species (Hu et al., 2011, Xu et al., 2013, Chiou et al., 2006, Du et al., 2018). Further studies demonstrated that induction of miR399 by Pi deficiency is significantly repressed in a loss-of-function Arabidopsis mutant, *phr1* (Bari et al., 2006), showing that *PHR1* acts as an upstream regulator of miR399 in Pi-deprivation response. Although *PHR1*, miR399 and *PHO2* define a conserved regulatory module that maintains Pi homeostasis in various plant species (Bari et al., 2006), it is possible that species-specific or even variety-specific Pi responses have also evolved.

Maize is not only an important crop for food and forage, but also a source of primary compounds for industrial innovation (Andorf et al., 2019, Godfrey, 2010). It has been widely cultivated in tropical and temperate soils worldwide, following the domestication and spread of its wild progenitor, teosinte (*Zea mays* ssp. *parviglumis*), originally grown in the subtropical environment of the Balsas River basin (Guerrero, México) (Matsuoka et al., 2002). Under cultivation, especially in acidic and alkaline soils, large amounts of Pi fertilizers are applied in maize fields; this allows maize plants to become well-established and to mature in a suitable timeframe, which improves the overall yield. However, despite these advantages, the long-term use of large amounts of fertilizers might result in the weakening of the Pi absorption system in plants during cultivation. Thus, research on Pi regulators with the ultimate aim of optimizing the acquisition and use of Pi by plants is needed.

Here, we report that overexpressing miR399 in maize resulted in typical Pi-toxicity phenotypes, as manifested by a premature aging syndrome in mature leaves after pollination. We also created *Zmpho2* mutants by CRISPR/Cas9 editing and found that knocking out the *ZmPHO2* gene, the miR399 target, consistently produced a premature aging phenotype similar to that of the transgenic miR399-overexpression lines. By performing transcriptome profiling and targeted metabolite measurements in miR399-overexpression and miR399-knockdown transgenic lines, we found that, in addition to affecting the Pi response, miR399 might mediate pathogen and insect defense in maize by regulating the expression of *Bx* genes in the benzoxazinoid biosynthesis pathway. We further demonstrated that an upstream regulator of the miR399-*ZmPHO2* module, INDETERMINATE1 (ID1), can inhibit the transcription of *ZmMIR399* genes and hence the accumulation of mature miR399, alleviate the repression of *ZmPHO2* by miR399, and ultimately maintain Pi homeostasis in maize. And *INDETERMINATE1 (ID1)* expression is unperturbed by low-Pi stress, suggesting that *ID1* acts as an autonomous Pi regulator in maize. In the current model, we conclude that mutation of *ID1* triggers the miR399-*ZmPHO2* pathway, a canonical and conserved Pi response module in plants, to promote Pi acquisition, providing the Pi supply for the vegetative gigantism of the *id1* mutant. We also showed that *ZmPHO2* was under selection during maize domestication and adaptation, resulting in a more sensitive response to phosphate starvation in temperate maize than tropical maize. We speculate that, during maize domestication or improvement, the Pi absorption system became suboptimal in cultivated maize due to prolonged and excessive Pi application in pursuit of improvements in yield. Our study establishes a regulatory connection between nutrient homeostasis and vegetative development in maize.

## Results

### Transgenic maize plants overexpressing miR399 display typical Pi-toxicity phenotypes and a premature aging syndrome after pollination

The miR399 family in maize contains a total of 10 members (i.e., miR399a-j) encoded by ten *ZmMIR399* genes in the genome and can be divided into six subgroups based on mature miRNA sequences (Zhang et al., 2009) (**Supplemental Figure S1, A and B**). To investigate the biological roles of miR399 in maize, we overexpressed two types of mature 21-nt miR399 sequences (miR399a/c/h and miR399e/i/j) in the same vector (**Supplemental Figure S2A**). Three independent transgenic lines (named miR399-OE#1, miR399-OE#2 and miR399-OE#3) were chosen for subsequent experiments after confirming miR399 overexpression by northern blot analysis (**Figure 1A**). The total phosphorus content was significantly higher in shoots of miR399-overexpressing transgene-positive plants (miR399-OEP#3) than in isogenic plants without the transgene, *i.e.* miR399 transgene-negative plants (miR399-OEN#3) derived from the same transformation experiment (**Figure 1B**), suggesting that overexpression of miR399 induces enhanced phosphate acquisition. In contrast, the phosphorus content in roots of transgene-positive and transgene-negative plants was not significantly different (**Figure 1B**), indicating that overexpression of miR399 improves phosphate mobilization from root to shoot. To investigate whether miR399 overexpression affects maize developmental and agronomic traits, we grew three independent miR399-overexpression lines in the field (Liaoning, 41.8° N, 123.4° E) along with their corresponding isogenic control plants. At 15 days after pollination (15-DAP), we observed markedly earlier leaf senescence in all three miR399 overexpression transgene-positive lines than in the corresponding transgene-negative lines **(Figure 1C; Supplemental Figure S3)**. Concomitantly, the miR399-overexpression transgenic lines showed significantly lower chlorophyll content across the entire mature leaf, from base to tip (**Figure 1D**).

**Figure 1:**
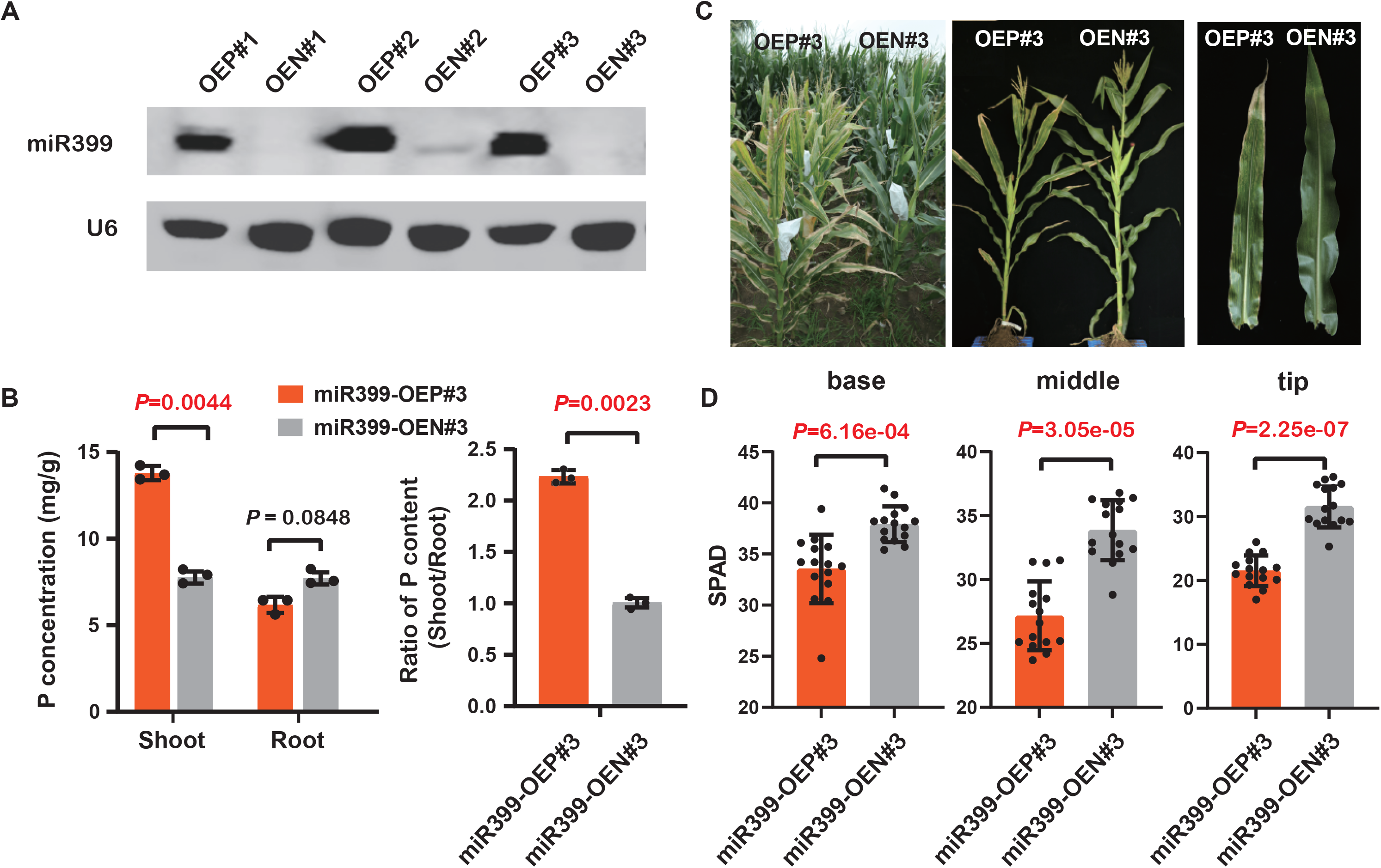
Phenotype of field-grown miR399-overexpressing transgenic plants. (A) Northern blot showing that miR399 over-accumulates as expected in three independent transgenic lines (named OEP#1 - OEP#3). A total of 10 µg total RNA from seedlings of each line was loaded per lane and the membrane was hybridized with a biotin-labeled miR399 probe. U6 served as a loading control. OEP and OEN represent transgene-positive and transgene-negative plants, respectively, derived from the same transformation event. (B) Total phosphorus (P) content in shoots and roots of miR399-overexpressing transgene-positive and transgene-negative lines. A representative transgenic strain, miR399-OE#3, was chosen and three biological replicates (n=3) for each genotype were prepared for total P concentration measurement. The data are shown as means ± SD; the *p*-values were determined by Student’s *t*-test. (C) Phosphate toxicity and premature aging phenotype of the miR399-overexpressing transgenic line (miR399-OE#3) grown in the field. A field scene and representative plants at 15 days after pollination are shown. (D) Quantification of total chlorophyll content (shown with SPAD values) in different parts (base, middle and tip) of mature ear leaves from OEP#3 and the corresponding transgene-negative control (OEN#3) after pollination. A total of 15 plants were measured for chlorophyll content for each genotype. The data are shown as means ± SD; the *p*-values were determined by Student’s *t*-test. SPAD represents total chlorophyll content, which was measured with the Konica Minolta SPAD 502 Plus instrument.

Using miRNA short tandem target mimicry (STTM) technology (Tang et al., 2012, Tang & Tang, 2013), we also generated miR399-knockdown transgenic plants (miR399-STTM) where the STTM RNA targets two miR399 species (corresponding to miR399a/c/h and miR399e/i/j), thereby covering the whole miR399 family **(Supplemental Figure S2B)**. The abundance of miR399 decreased in two independent miR399-STTM transgene-positive lines (miR399-STTMP#1 and miR399-STTMP#2) compared to their isogenic siblings without the transgene (miR399-STTMN#1 and miR399-STTMN#2), confirming the successful knockdown of endogenous miR399 **(Supplemental Figure S4A)**. Phosphorus accumulation in shoots was not significantly different between miR399-STTMP#2 and miR399-STTMN#2 lines **(Supplemental Figure S4B)**. The lack of a change in phosphorus content might be due to the naturally low levels of miR399 at the seedling stage under normal growth conditions, which would minimize any differences in miR399 levels between miR399-STTMP and miR399-STTMN lines. Homozygous miR399-STTMP transgenic lines were grown in the field (Liaoning, 41.8° N, 123.4° E) for phenotype observation along with their corresponding isogenic negative controls (miR399-STTMN#1 and miR399-STTMN#2). No visible developmental or morphological differences between the transgene-positive and the corresponding transgene-negative lines were observed **(Supplemental Figure S4C; Supplemental Figure S5)**, implying that a minor reduction in miR399 levels does not affect maize development under normal conditions. Collectively, these results indicate that miR399 functions as a positive regulator of phosphorus absorption and that overexpression of miR399 induces a premature aging syndrome after pollination in maize.

### Knocking out *ZmPHO2*, a miR399 target, results in premature senility

*PHO2,* which encodes a UBC enzyme, has been experimentally confirmed to be a target of miR399 in various species, including Arabidopsis, rice, barley, soybean and maize (Hackenberg et al., 2013, Fujii et al., 2005, Hu et al., 2011, Xu et al., 2013, Du et al., 2018, Bari et al., 2006, Chiou et al., 2006, Pant et al., 2008). In maize, miR399 was found to direct the cleavage of *ZmPHO2* mRNA, which contains six tandem miR399-binding sites in its 5’ UTR (**Figure 2A**), by transient expression and 5’-RACE assays (Du et al., 2018). We examined *ZmPHO2* expression by qRT-PCR in miR399-OE#3 and miR399-STTM#2 transgenic lines and found the expected expression changes with significant downregulation and upregulation in miR399-OE and miR399-STTM transgenic lines, respectively (**Figure 2B**), further confirming that *ZmPHO2* is an authentic target of miR399. However, so far, there is no genetic evidence for the function of *ZmPHO2* in the regulation of phosphate homeostasis and plant development in maize. To examine the function of *ZmPHO2*, we sought to knock out *ZmPHO2* in the maize inbred line ZZC01 using CRISPR/Cas9-mediated mutagenesis (Doudna & Charpentier, 2014, Belhaj et al., 2015, Ma et al., 2015). We designed two single guide RNAs to target the coding region immediately after the translation start site within the first exon of *ZmPHO2* **(Figure 2, A and C)**. Genotyping and sequence analyses identified three homozygous mutants harboring fragment insertions and/or deletions around the guide RNA sites (**Figure 2C**). The three homozygous, potentially null mutants consistently showed a typical Pi-toxicity phenotype and exhibited premature aging (**Figure 2D**). The phenotypes were more severe than those of the miR399 OE transgenic lines such that very few seeds could be harvested. Similar phenotypes were also observed in *pho2* mutants of rice and Arabidopsis, in which the over-accumulated Pi in their leaves resulted in symptoms of Pi toxicity and premature senescence, when grown in Pi-sufficient soil (Delhaize & Randall, 1995, Wang et al., 2009).

**Figure 2:**
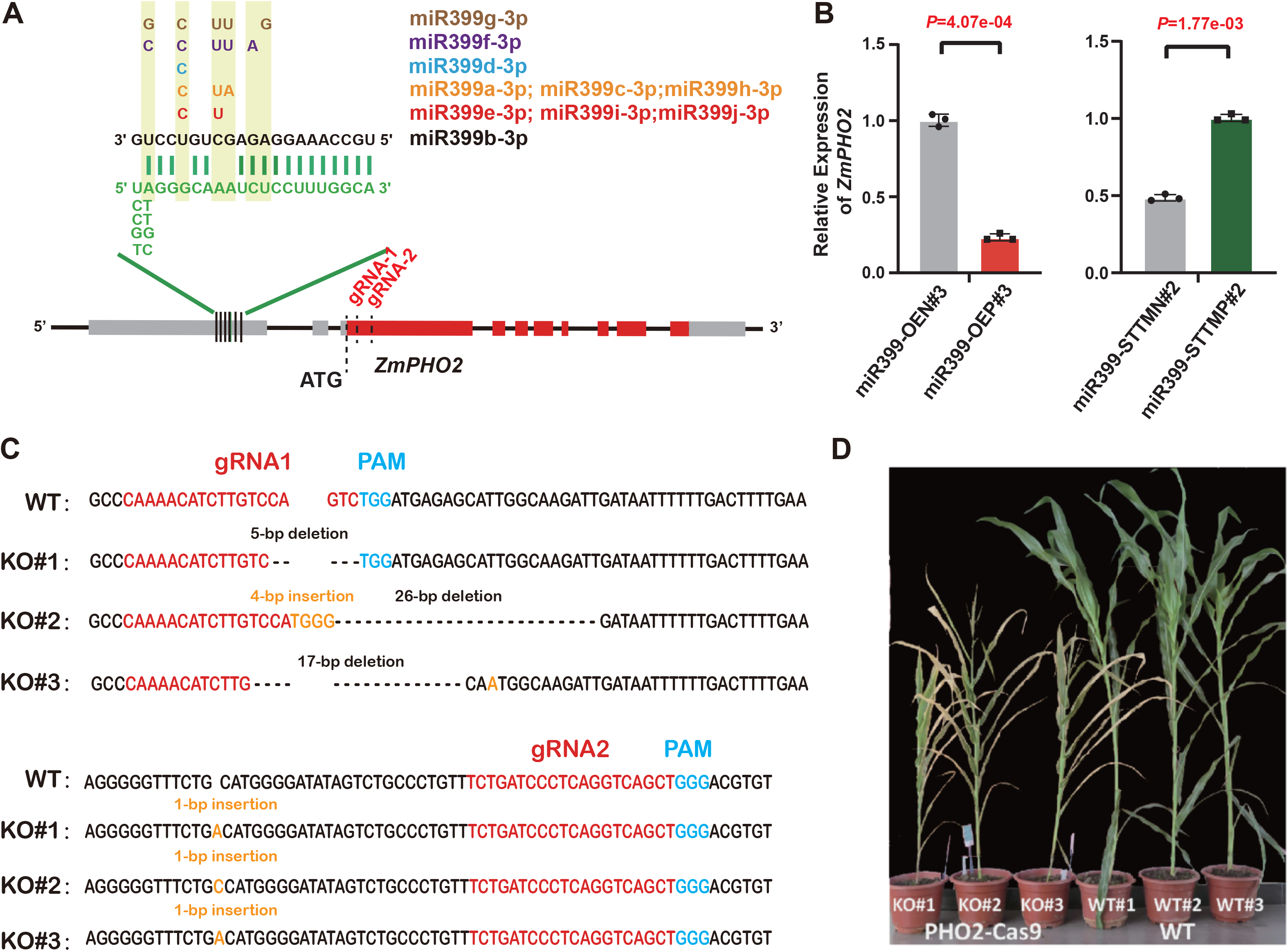
CRISPR/Cas9-mediated target mutagenesis of *ZmPHO2* causes premature senescence in maize. (A) Diagram of *ZmPHO2* showing the base complementarity of the miR399 target sites and mature miR399 members. Six predicted miR399 target sites (four of them were experimentally validated by Du et al. (2018)) are located in the 5’ UTR region. Grey and red boxes in the diagram represent UTR and coding regions of *ZmPHO2*, respectively. gRNA-1 and gRNA-2 represent the two guide RNAs (orange lines) for CRISPR/Cas9 editing. (B) *ZmPHO2* expression evaluated by qRT-PCR in miR399-OE and miR399-STTM transgene-positive lines and the corresponding negative controls. The *p*-values are calculated by Student *t*-test. The y-axis represents the expression level relative to the *ZmUbiqutin* gene. (C) Sequences of three homozygous knockout lines with insertions/deletions that truncate the *ZmPHO2* open reading frame (named KO#1, KO#2, and KO#3). The wild-type (WT) sequence is shown at the top. Guide RNA and protospacer-adjacent motif (PAM) sequences are in red and blue text, respectively. The deletions are indicated by dashes, while the insertion was showed with orange nucleotides. (D) Gross morphology of three *ZmPHO2* knockout and three wild type lines.

### miR399 might mediate pathogen and insect defense in maize, in addition to affecting Pi response

To characterize the gene network(s) through which miR399 might impact maize development and physiology, we carried out a comparative transcriptomics analysis of mature leaves (V5 stage) between homozygous miR399-OEP#3 and miR399-STTMP#2, and their corresponding transgene-negative lines (miR399-OEN#3 and miR399-STTMN#2, respectively) with three reproducible replicates **(Supplemental Figure S6)**. By comparison with the corresponding transgene-negative lines, we identified 2,407 and 1,183 differentially expressed genes (DEGs) in miR399-OE#3 and miR399-STTM#2, respectively (Fold Change >=1.5 & FDR <=0.05; **Figure 3A; Supplemental Dataset S1; Supplemental Dataset S2)**. Of these, 903 genes were defined as hypo-DEGs in miR399-OE plants, with significantly lower expression levels observed in miR399-OEP#3 than in miR399-OEN#3 (**Figure 3A**). Similarly, 629 hyper-DEGs were identified in miR399 STTM plants, with significantly higher expression detected in miR399-STTMP#2 than in miR399-STTMN#2 (**Figure 3B**). Overlap analysis between the hypo-DEGs in miR399 OE and hyper-DEGs in miR399 STTM identified a small subset of highly confident genes (101 genes; **Figure 3C**), whose expression was negatively correlated with miR399 levels. Annotation of these genes revealed a number of Pi-related genes, including *ZmPHO2* **(Figure 3D; Supplemental Dataset S2)**. Pathway enrichment analysis of these miR399-associated genes in reference to the Gene Ontology (GO) and Kyoto Encyclopedia of Genes and Genomes (KEGG) databases showed that the term “benzoxazinoid biosynthesis” was the most strongly enriched (**Figure 3E**).

**Figure 3:**
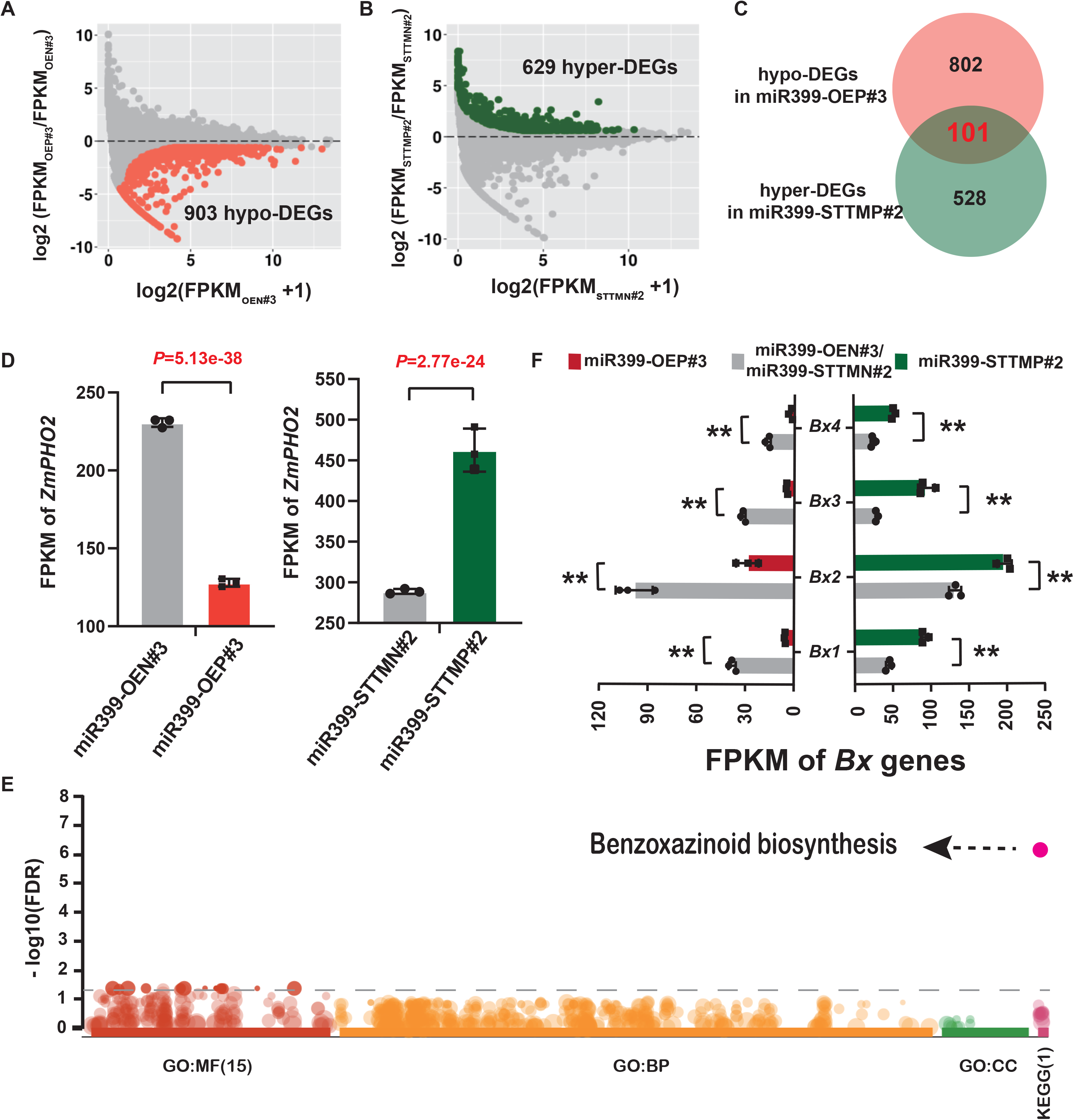
RNA-seq analysis reveals that miR399 might mediate pathogen and insect resistance in maize, in addition to affecting Pi response genes. (A) Differentially expressed genes (DEGs) in the miR399-OE transgene-positive line (miR399-OEP#3) compared with the transgene-negative line (miR399-OEN#3). A total of 903 downregulated genes (hypo-DEGs) were detected in miR399-OEP#3. (B) DEGs in the miR399-STTM transgene-positive line (miR399-STTMP#2) compared with the transgene-negative line (miR399-STTMN#2). A total of 629 upregulated genes (hyper-DEGs) were detected in miR399-STTMN#2. (C) Overlap of the hypo-DEGs identified in miR399-OEP#3 and hyper-DEGs in miR399-STTMP#2. (D) *ZmPHO2* expression in miR399-OE and miR399-STTM transgene-positive lines (miR399-OEP#3 and miR399-STTMP#2) and the corresponding transgene-negative lines (miR399-OEN#3 and miR399-STTMN#2). *p*-values were calculated by DESeq2 after FDR multiple correction. (E) Gene ontology (GO) and KEGG pathway enrichment analysis of the overlapping DEGs (the 101 genes indicated in C). Benzoxazinoid biosynthesis was the most strikingly enriched pathway. BP, MF and CC represent Biological Process, Molecular Function, and Cellular Component groups of GO, respectively. The total number of categories in the GO or KEGG group is indicated in parentheses. (F) Expression of several key *Bx* genes (*Bx1-Bx4*) involved in benzoxazinoid biosynthesis pathway in the miR399-OE and miR399-STTM transgene-positive lines and the corresponding transgene-negative lines. **, *p*-value <=0.01.

Benzoxazinoids are a class of indole-derived protective and allelopathic secondary metabolites that are involved in anti-fungal and insect defense responses in plants (Frey et al., 2009). 2,4-dihydroxy-1,4-benzoxazin-3-one (DIBOA) and its 7-methoxy analog 2,4-dihydroxy-7-methoxy-1,4-benzoxazin-3-one (DIMBOA) are the predominant benzoxazinoids in plants (Frey et al., 2009, Frey et al., 1997) and these compounds have been found in many plants including maize, wheat and rye (*Secale cereale*) (Frey et al., 2009). The benzoxazinoid biosynthetic pathway is well-characterized in maize **(Supplemental Figure S7A)**. Previously, metabolic profiling revealed that levels of both DIBOA-glucoside (DIBOA-Glc) and DIMBOA-glucoside (DIMBOA-Glc), the storage forms of DIBOA and DIMBOA, are markedly decreased in Pi-resistant maize lines, which are able to absorb Pi more efficiently under conditions of Pi deficiency (Luo et al., 2019). In our present study, the *Bx* genes (*Bx1-Bx9*) involved in benzoxazinoid biosynthesis were significantly downregulated in miR399-OE transgene-positive lines, as shown by the RNA-seq data and qRT-PCR validation **(Figure 3F; Supplemental Figure S7B; Supplemental Figure S8)**, which should result in a reduced accumulation of benzoxazinoid compounds. To test this, we performed a targeted high performance liquid chromatography (HPLC) assay with DIBOA and DIMBOA as standards. The results showed that levels of both DIBOA and DIMBOA were significantly reduced in miR399-OE plants, as hypothesized **(Supplemental Figure S9A)**. In contrast, the expression levels of most *Bx* genes tended to be upregulated and the metabolites (DIBOA and DIMBOA) accumulated to higher levels in miR399-STTM plants **(Figure 3F; Supplemental Figure S8; Supplemental Figure S9B)**. Collectively, these results support the conclusion that miR399 might impact pathogen and insect defense in maize, in addition to its influences on the Pi response. However, how the expression of the *Bx* genes is modulated in miR399-OE or miR399-STTM transgenic lines is still an open question. A similar scenario has been observed in rice, where Pi accumulation induced by overexpressing *OsMIR399f* or high levels of Pi fertilization results in enhanced susceptibility to infection by the blast fungus *Magnaporthe oryzae* (Campos-Soriano et al., 2020), demonstrating that miR399 functions as a negative regulator of plant immunity, possibly through an indirect mechanism.

### Mutation of ID1 promotes miR399 expression and facilitates Pi accumulation in maize

The ID1 protein, a monocotyledon-specific zinc-finger transcription factor, plays an essential role in regulating the transition to flowering in maize. Loss-of-function *id1* mutants experience a prolonged period of vegetative growth and concomitantly extremely delayed flowering, ultimately resulting in vegetative gigantism (Colasanti & Sundaresan, 2000, Singleton, 1946, Colasanti et al., 1998). Although the *id1* mutation severely alters the ability of maize to undergo the transition to reproductive growth, homozygous *id1* plants are virtually identical in growth rate and morphology to normal maize plants until the point of floral transition (Coneva et al., 2007, Colasanti et al., 1998, Singleton, 1946). Transcriptome profiling revealed that miR399 accumulated to abnormally high levels in both mature and immature leaves of the *id1-m1* mutant at the floral transition stage (Minow et al., 2018). By re-analyzing the small RNA-seq data from this study (Minow et al., 2018), we confirmed that mature members of the miR399 family (including miR399a/c/h, miR399e/i/j, miR399b and miR399f) were significantly upregulated in mature leaves of the *id1-m1* mutant **(Figure 4A; Supplemental Figure S10)**. In immature leaves, miR399e/i/j also accumulated to higher levels in the *id1-m1* mutant **(Supplemental Figure S10)**, while other members were slightly upregulated or unchanged. Using the corresponding RNA-seq data (Minow et al., 2018), we found that *ZmPHO2* was significantly downregulated in the *id1-m1* mutant **(Supplemental Figure S11)**. Considering the nutritional requirements for vigorous vegetative growth and extremely delayed flowering, we speculate that the upregulation of miR399 in the *id1* mutant might help to provide more Pi to sustain the extra plant growth. This hypothesis prompted us to examine the relationship between ID1 and the miR399-*ZmPHO2* regulatory module.

**Figure 4:**
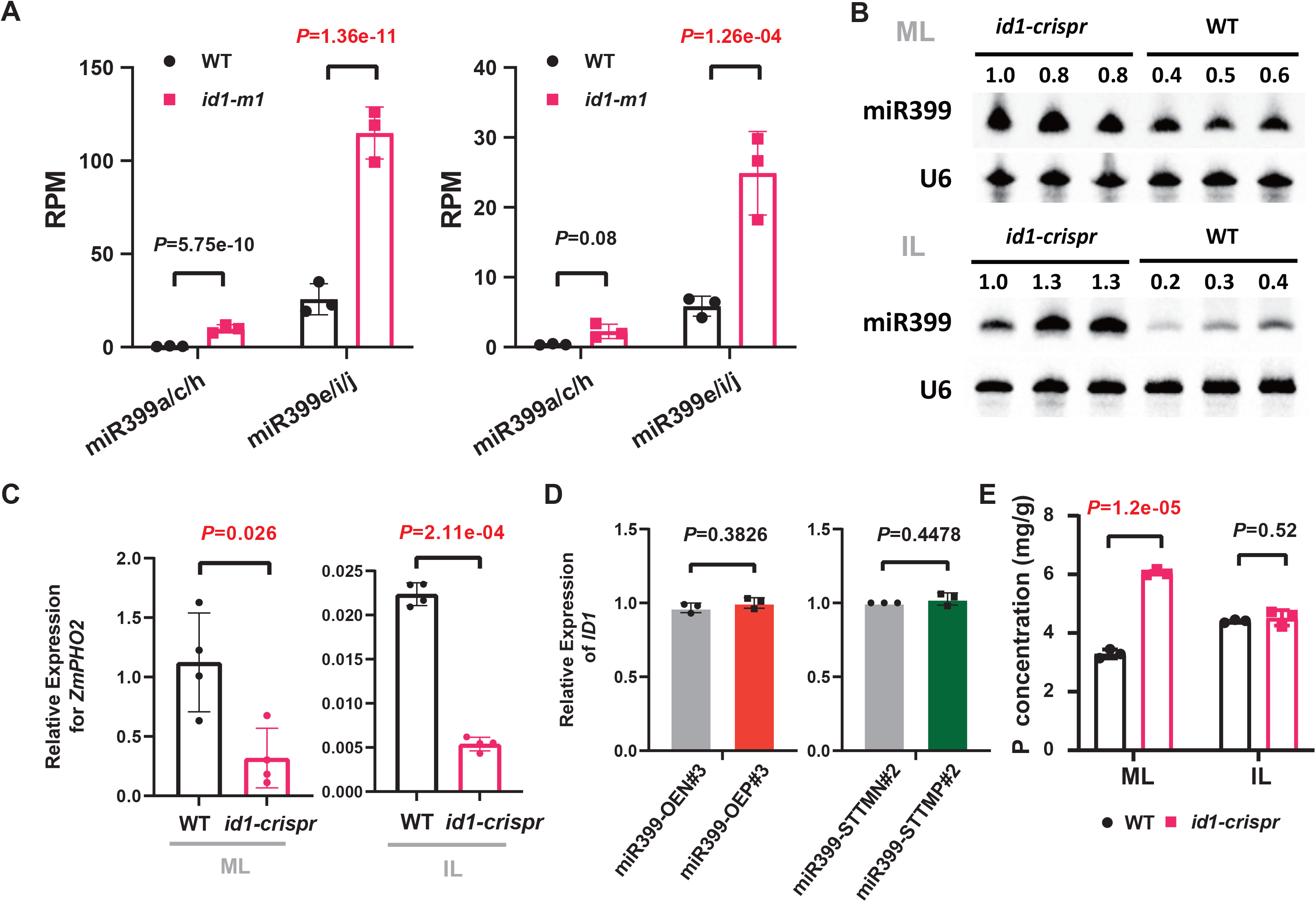
Mutation of *ID1* promotes miR399 accumulation and facilitates Pi accumulation in maize. (A) Abundance of miR399a/c/h and miR399e/i/j in the *id1-m1* mutant and wild-type (WT) based on small RNA-seq data. The left and right panels represent results of small RNA-seq in mature leaves and immature leaves, respectively. The y-axis indicates miRNA abundance (RPM, reads per million). The data are shown as means ± SD. *p*-values were calculated by DESeq2 after FDR multiple correction. (B) Northern blot to detect miR399 abundance in mature leaves (ML) and immature leaves (IL) of the *id1-crispr* mutant and the wild type (WT). U6 served as an internal control. The numbers indicate the abundance of miR399 relative to the first lane. (C) *ZmPHO2* expression levels in mature leaves (ML) and immature leaves (IL) of the *id1-crispr* mutant (n = 3). The y-axis represents the expression level relative to that of the *ZmUbiqutin* gene. (D) *ID1* expression levels in miR399-OE (OE#3) transgenic lines and miR399-STTM (STTM#2) transgenic lines (n = 3). The y-axis represents the expression level normalized to that of the *ZmUbiqutin* gene. (E) Total phosphorus (P) content in mature leaves (ML) and immature leaves (IL) of the *id1-crispr* mutant and wild type (WT) at the transition stage. The data in panels C-E are presented as means ± SD. Student’s *t*-test was employed to calculate the *p*-value.

To independently confirm the small RNA-seq results, we first created a novel *id1* mutant by CRISPR/Cas9 gene editing technology (Ma et al., 2015, Belhaj et al., 2015) and obtained an edited plant with a large deletion in the *ID1* coding region **(Supplemental Figure S12A).** By genotyping and sequencing the targeted genomic region in T2 populations derived from a T1 heterozygous mutant, we identified a homozygous mutant without the CRISPR/Cas9 transgene. The homozygous mutant, which was named *id1-crispr*, showed lower expression of *ID1* **(Supplemental Figure S12B)** and, as expected, vigorous vegetative growth and extremely delayed flowering in the field **(Supplemental Figure S12, C and D)**. These molecular and phenotypic characteristics indicated that we had successfully obtained an *id1* mutant by CRISPR/Cas9 technology. By RNA gel blot analysis, we confirmed that miR399 levels were significantly increased in mature and immature leaves at the floral transition stage in the *id1-crispr* mutant (**Figure 4B**). In addition, consistent with the reported RNA-seq data **(Supplemental Figure S11)** (Minow et al., 2018), the expression of *ZmPHO2* was confirmed by qRT-PCR to be significantly downregulated in the *id1-crispr* mutant (**Figure 4C**). We further examined the expression of *ID1* in miR399-OE and miR399-STTM transgenic lines by RNA-seq and qRT-PCR, and did not detect any significant difference from wild type **(Figure 4D; Supplemental Figure S13)**, which supports the notion that *ID1* is genetically upstream of the miR399-*ZmPHO2* module. Next, we measured the total phosphorus content in *id1-crispr* mutant and found two-fold higher phosphorus levels in mature leaves compared to wild type (**Figure 4E**). In other tissues, including immature leaf, root, leaf sheath and stem, no significant phosphorus content changes were observed **(Supplemental Figure S14)**. Altogether, these observations show that loss of ID1 function facilitates Pi accumulation in mature leaves, perhaps via the miR399-*ZmPHO2* regulatory module. Increased Pi accumulation might provide an adequate nutritional reserve for the vigorous vegetative growth of the *id1* mutant.

### ID1 directly binds to the promoters of *ZmMIR399c* and *ZmMIR399j* to repress their expression

We determined the levels of the primary transcripts (pri-miR399s) derived from the *ZmMIR399a, ZmMIR399c, ZmMIR399h, ZmMIR399e, ZmMIR399i* and *ZmMIR399j* genes, which are responsible for the production of miR399a/c/h and miR399e/i/j. We found that, in both mature and immature leaves of the *id1-crispr* mutant, *ZmMIR399c* and *ZmMIR399j* were markedly upregulated, while the expression of other *ZmMIR399* genes was slightly upregulated or unchanged (**Figure 5A**), indicating that the enhanced accumulation of miR399a/c/h and miR399e/i/j in the *id1-crispr* mutant was mainly due to *ZmMIR399c* and *ZmMIR399j* expression, respectively. Next, we examined whether ID1 could regulate the transcription of *ZmMIR399c* and *ZmMIR399j* using a dual-luciferase (LUC) reporter assay, in which the coding sequence of *ID1*, driven by the 35S promoter (*p35S::ID1*), was used as the effector and *LUC* driven by either the *ZmMIR399c* or the *ZmMIR399j* promoter (*pro-MIR399c::LUC and pro-MIR399j::LUC*), was used as the reporter (**Figure 5B**). We found that the *pro-MIR399c::LUC* and *pro-MIR399j::LUC* reporters were significantly repressed in maize protoplasts when the *p35S::ID1* effector was co-expressed (**Figure 5B**).

**Figure 5:**
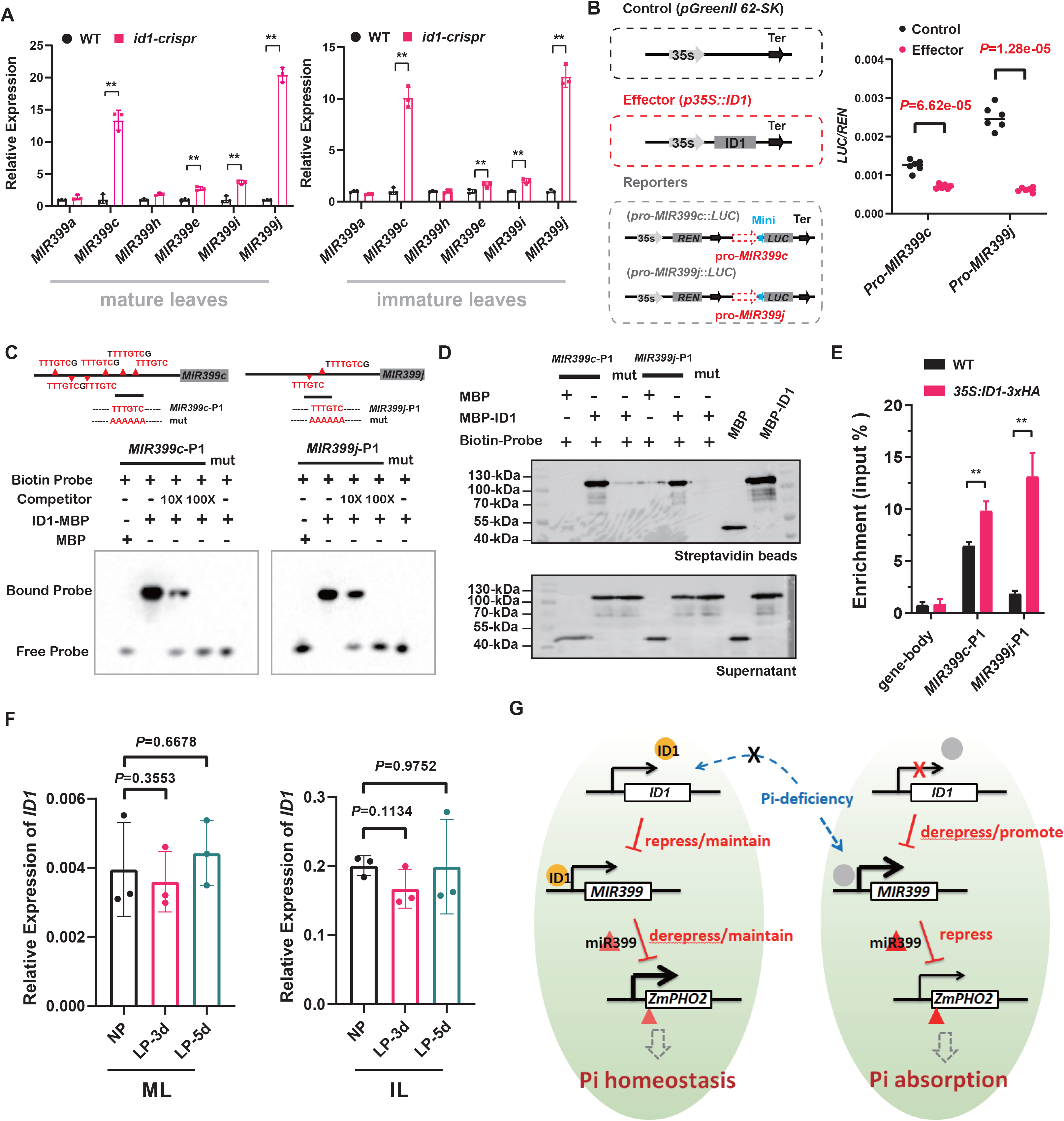
ID1 functions as an upstream regulator by binding directly to the promoter regions of *MIR399c* and *MIR399j*. (A) Expression comparison by qRT-PCR of six primary miR399 precursors (*MIR399a, c, h, e, i, j*) in mature leaves and immature leaves of the *id1-crispr* mutant and wild type (WT). The y-axis represents expression levels relative to the *ZmUbiqutin* gene. The data are shown as means ± SD; the *p*-values were determined by Student’s *t*-test. (B) Schematic diagram showing the effector and reporter constructs used in the transient transcription activity assays. *REN*: Renilla luciferase; *LUC*: firefly luciferase; Mini: 35S minimal promoter, which confers a basal level of *LUC* expression. Co-expressing the effector (*p35S::ID1*) with two reporters (*pro-MIR399c::LUC* and *pro-MIR399j::LUC*) gave a significantly lower LUC activity. The data are presented as means ± SD based on the technical replicates (n = 6); Student’s *t*-test was used to calculate the *p*-values. The experiment was repeated three times independently and a representative result is shown. (C) EMSA assay showing the binding of ID1 to the promoters of *MIR399c* and *MIR399j*. A diagram showing the predicted binding motif distribution for the ID1 transcription factor along the promoter regions of *MIR399c* and *MIR399j* is shown at the top. The black lines show the biotin-labeled probes used in EMSA and *in vitro* DNA pull-down experiments. The bottom panels show the binding of ID1 protein to the predicted motifs in the promoters of *MIR399c* and *MIR399j*. “mut” represents the mutated biotin-probes *MIR399c*-p1 and *MIR399j*-p1 **(Supplemental Dataset 4)**. (D) DNA pull-down analysis showing the binding of ID1 to *MIR399c* and *MIR399j* promoters at the predicted motifs. Biotin-labeled probes, including *MIR399c*-P1 and *MIR399j*-P1, were used to pull-down recombinant MBP (control) and MBP-ID1. Results were analyzed by western blot using anti-MBP antibody. The supernatant was used as an internal control. (E) Chromatin immunoprecipitation (ChIP) to show the association of ID1 with the promoters of *MIR399c* and *MIR399j in vivo*. Chromatin from the leaves of *35S::ID1-3*x*HA* transgenic or wild-type (WT) plants was immunoprecipitated with anti-HA Trap. The enrichment (Input %) of the two promoter fragments (*MIR399c*-P1 and *MIR399j*-P1 shown in (C)) for each genotype was determined by quantitative PCR. Asterisks indicate significant difference between the *35S::ID1-3xHA* transgenic line and wild type (WT) (**, *p*-value < =0.01; Student’s *t-*test). (F) Relative expression of *ID1* under low-Pi stress conditions. NP, normal Pi condition; LP-3d, 3 days under low-Pi conditions; LP-5d, 5 days under low-Pi conditions; ML, mature leaves; IL, immature leaves. qRT-PCR was performed with three biological replicates and *ZmUbiqutin* was used as a reference gene for normalization. The data are shown as means ± SD, and the *p*-values were determined by Student’s *t*-test. (G) Proposed model for the function of ID1 as a Pi homeostasis regulator. In the absence of Pi stress, ID1 regulates the expression of *MIR399* genes to maintain *in vivo* Pi homeostasis and normal metabolism. When ID1 is mutated, de-repression of *MIR399c* and *MIR399j* promotes over-accumulation of mature miR399, and elevated miR399 levels inhibit the expression of *ZmPHO2*, resulting in enhanced Pi acquisition and transport. As a result, adequate Pi nutrition is stored in leaves to support the extra growth of the *id1* mutant. The size of the black arrows indicates the gene expression level. Arrows denote positive regulation and blunt-ended lines indicate inhibition.

The transcription factor ID1 has been shown to selectively bind to an 11-bp consensus element by *in vitro* DNA selection and amplification binding (SAAB) assays and DNA binding experiments (Kozaki et al., 2004). Transient expression in *Nicotiana benthamiana* showed that ID1 was mainly localized in the nucleoplasm **(Supplemental Figure S15)**, in line with its potential regulatory function as a transcription factor. Sequence analysis of the 2-kb regions upstream of the annotated 5’ ends of *ZmMIR399c* and *ZmMIR399j* revealed the presence of six and two copies, respectively, of a typical ID1-binding motif **(Figure 5C; Supplemental Figure S16)**. To test whether ID1 can bind directly to the promoter regions, we purified a maltose-binding protein tagged version of ID1 (MBP-ID1) protein in *E. coli* **(Supplemental Figure S17)**, and performed electrophoretic mobility shift assay (EMSA) and *in vitro* DNA pull-down experiments by incubating biotin-labeled probes with the recombinant MBP-ID1 protein. The results indicated that ID1 can bind to DNA fragments containing the motifs, while corresponding non-biotin-labeled competitors neutralized the binding activity **(Figure 5, C and D)**. Mutation of the motif sequences completely abolished ID1 binding **(Figure 5, C and D)**. Lastly, we utilized the *35S::ID1-3*x*HA* transgenic line **(Supplemental Figure S18)** to conduct a chromatin immunoprecipitation (ChIP) experiment in leaf tissue. Immunoprecipitation with an HA antibody showed a specific enrichment of ID1-3xHA in the promoter regions of *ZmMIR399c* and *ZmMIR399j* that encompass the probe sequences **(Figure 5, C and E)**. Taken together, these results demonstrate that ID1 directly binds to the promoter regions of *ZmMIR399c* and *ZmMIR399j* and represses their transcription.

We further investigated the response of *ID1* expression to low-Pi stress by growing the B73 inbred line in hydroponic solutions containing normal Pi (NP; 250 μM KH_2_PO_4_) and low Pi (LP; 5μM KH_2_PO_4_). Low-Pi stress treatment was implemented for three or five days prior to the reproductive transition stage when *ID1* is specifically expressed. Mature and immature leaves were collected and used for RNA isolation and expression analysis. We first confirmed that the levels of the Pi-responsive miR399 and the long non-coding RNA *PILNCR1* were elevated under low-Pi conditions **(Supplemental Figure S19)**, indicating that the Pi treatment was effective. However, we did not detect expression changes for the *ID1* gene under low-Pi conditions (**Figure 5F**), suggesting that ID1 is not a Pi-responsive, but an autonomous regulator in maize.

Based on all of the above observations, we propose that loss of *ID1* triggers the miR399-*ZmPHO2* pathway, which enhances Pi acquisition and transport, and finally leads to higher Pi levels that are able to support the vegetative gigantism of the *id1* mutant (**Figure 5G**). This contrasts with wild-type plants, where ID1 downregulates miR399 levels by repressing the transcription of *ZmMIR399c and ZmMIR399j*, which in turn maintains Pi homeostasis (**Figure 5G**), highlighting that ID1 is an autonomous Pi regulator upstream of the miR399-*ZmPHO2* signaling pathway.

### *ZmPHO2* was under selection during maize domestication

Crop domestication has not only been shaped by artificial selection but also by natural selection pressures (Martín-Robles et al., 2019). Natural selection under modern agricultural conditions, which differ from wild habitats in the availability of resources, might have led to adaptations of crops in both external morphology and internal metabolism. Cultivated maize originated in the tropical environment of southwestern Mexico and has since spread widely to temperate zones having topsoil properties that differ from those of tropical areas. In tropical and subtropical regions, such as the Cerrado region in Brazil, acidic soils with low Pi availability are prevalent (Goedert, 1983, Aguirre-Liguori et al., 2019). Under these conditions, without human interference, the ancestral maize may have needed a more efficient Pi absorption system to ensure its survival. While in humid temperate regions, Pi seems to be concentrated at the surface, a considerable part of it in organic compounds (Lynch & Brown, 2001). Under these conditions, cultivated maize can have better access to Pi. Furthermore, the use of fertilizer, which tends to be over-applied by farmers to guarantee adequate levels of crop yield and quality, makes Pi easily available. Thus, both natural and artificial selection pressures might have promoted changes in the Pi absorption/transport system as maize was domesticated from teosinte. We speculate that *ZmPHO2*, as a major phosphate regulator, might have been targeted by selection as maize spread to temperate zones.

To test this hypothesis, we first analyzed the nucleotide diversity around ten *MIR399* genes and *ZmPHO2* using the third-generation *Zea mays* haplotype map (HapMap 3), which includes ultra-high-density single nucleotide polymorphism (SNP) data for various maize accessions (Bukowski et al., 2018). We observed that the regions around *ZmMIR399c* and *ZmMIR399j* showed a slight decrease in nucleotide diversity in maize or landraces compared to the teosinte group **(Supplemental Figure S20)**. Notably, the 5’ UTR of *ZmPHO2*, in which the miR399 binding sites are located, exhibited a clear reduction in nucleotide diversity in maize compared to landraces or teosinte, with the maize group retaining only 29% of the genetic diversity of the teosinte group (**Figure 6A**). We confirmed this observation by re-sequencing a ∼1.6-kb 5’ UTR region in 26 diverse maize inbred lines and 14 teosinte accessions (*Zea mays* ssp. *parviglumis*) **(Supplemental Dataset S3).** The results indicated that maize on average (π = 0.00266) retained 33.7% of the nucleotide diversity of teosinte (π = 0.00788; **Figure 6B**), with two specific regions (∼1.2-kb and ∼400-bp upstream of the ATG) showing very low nucleotide diversity in maize than in teosinte **(Supplemental Figure 21)**. Coalescent simulations that incorporate the domestication bottleneck (Tian et al., 2009, Eyre-Walker et al., 1998) detected a significant deviation from expectation under a neutral domestication bottleneck for this re-sequenced region **(P < 0.5;** **Figure 6B**), indicating that the 5’ UTR of *ZmPHO2* was affected by selection. Although the miR399 binding sites did not show any sequence differences among the re-sequenced maize and teosinte accessions, the SNPs exist around the miR399 binding sites, which might affect the accessibility of miR399 and thus alter the expression of *ZmPHO2*. With our previously published eQTL mapping results using a maize-teosinte recombinant inbred line (RIL) population (Wang et al., 2018), we detected a strong *cis*-eQTL for *ZmPHO2*, with the maize allele showing a significantly higher expression level than the teosinte allele (**Figure 6C**). We developed near-isogenic lines (NILs) across the *ZmPHO2* region from the same RIL population, and validated that the expression of *ZmPHO2* was significantly higher in *ZmPHO2*-NIL^maize^ than in *ZmPHO2*-NIL^teosinte^ (**Figure 6D**). In agreement with a role for *ZmPHO2* as a negative regulator of Pi uptake, *ZmPHO2*-NIL^maize^ lines exhibited a significantly lower total Pi level than *ZmPHO2*-NIL^teosinte^ lines (**Figure 6E**). These results indicate that *ZmPHO2* may have been a target of selection during maize domestication, leading to weaker Pi absorption in cultivated maize. We predicted the RNA secondary structures of the 5’ UTR region containing the miR399 binding sites (including 100-bp upstream of the first binding site and 100-bp downstream of the sixth binding site) from *ZmPHO2*-NIL^maize^ and *ZmPHO2*-NIL^teosinte^. Three SNPs were present in this region between *ZmPHO2*-NIL^maize^ and *ZmPHO2*-NIL^teosinte^, and the RNA secondary structures were predicted to be greatly affected by the SNPs **(Supplemental Figure 22)**. The fourth miR399 bind site appeared to be more exposed in *ZmPHO2*-NIL^teosinte^ **(Supplemental Figure 22)**, which might be more accessible to mature miR399, resulting in the lower expression of *ZmPHO2*. We further examined the difference in expression of *ZmPHO2* between temperate and tropical maize inbred lines using published RNA-seq data of the association panel (Kremling et al., 2018). The analysis showed that temperate lines tended to have higher *ZmPHO2* expression levels than tropical lines in root tissues (**Figure 6F**). In addition, a recent study evaluated low-Pi tolerance in another maize association panel, which included different sources of maize inbred lines (Fang & Luo, 2019). Using the classification criteria of that study, we found a greater percentage of low-Pi sensitive lines and a lower percentage of low-Pi resistant lines in the temperate maize subgroup than in the tropical maize subgroup (**Figure 6G**), implying that the Pi absorption system might be failing in domesticated maize, perhaps owing to the overuse of Pi fertilization. Taken together, the above results suggest that *ZmPHO2* expression was affected by domestication and may therefore have played a role in the adaptation of maize to temperate soils.

**Figure 6:**
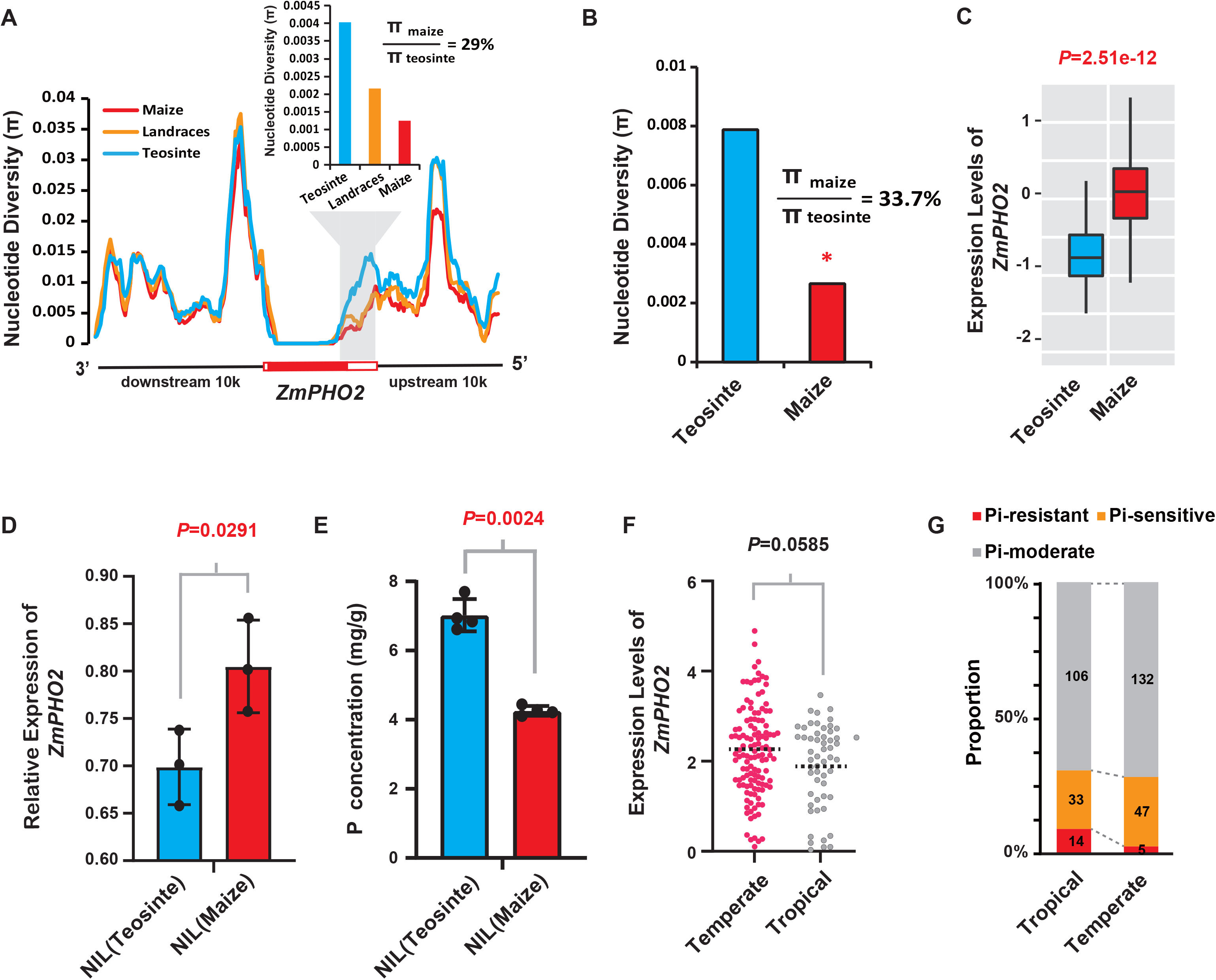
Evidence of selection at *ZmPHO2*. (A) The 5’ regulatory region of *ZmPHO2* exhibits low nucleotide diversity (*π*) in maize relative to teosinte, analyzed with maize HapMap v3 SNP data. The red, blue and orange curves represent the nucleotide diversity of maize, teosinte and landraces around the analyzed genes, respectively. The white and red rectangles on the x-axis indicate the UTR and CDS regions of the *ZmPHO2* gene. (B) Re-sequencing 26 maize inbreds and 14 teosinte accessions indicated that maize on average retained 33.7% nucleotide diversity of teosinte in the re-sequenced 5’ UTR region. * P <0.05 from the coalescent simulations. (C) *ZmPHO2* expression levels in maize and teosinte, based on data from Wang et al. (2018). (D) *ZmPHO2* expression levels in *ZmPHO2-NIL^maize^* and *ZmPHO2-NIL^teosinte^*examined by qRT-PCR. The y-axis represents the expression level normalized to that of the *ZmUbiqutin* gene. The data are presented as means ± SD. Student’s *t*-test was employed to calculate the *p*-value. (E) Comparison of total phosphorus (P) content between *ZmPHO2-NIL^maize^* and *ZmPHO2-NIL^teosinte^*. The data are presented as means ± SD. Student’s *t*-test was employed to calculate the *p*-value. (F) *ZmPHO2* expression levels in the root tissues of tropical and temperate inbred maize lines, based on data from (Kremling et al., 2018). *p*-values were calculated by Student’s *t*-test (G) Statistical survey of Pi-resistant, Pi-sensitive and Pi-moderate lines in tropical and temperate maize groups. The data are from (Luo et al., 2019).

## Discussion

### ID1, as an upstream repressor, might regulate the miR399*-ZmPHO2* module independently of known regulators

Several regulators involved in maintaining Pi homeostasis have been identified in plants (Chen et al., 2007, Devaiah et al., 2009, Devaiah et al., 2007b, Bari et al., 2006, Zhou et al., 2008, Wang et al., 2013b, Rubio et al., 2001). *AtPHR1*, which encodes a MYB-type transcription factor, was the first regulator demonstrated to mediate the Pi-starvation response; it plays a central role in transcriptional regulation of a large subset of Pi-deficiency-responsive genes, including *ZmMIR399* (Bari et al., 2006). In the promoter regions of *ZmMIR399* genes, AtPHR1 binds the predicted PHR1-specific binding sequence (P1BS) motifs, resulting in positive regulation of miR399 accumulation. In a similar manner, *PHR1* orthologs have been reported to act as transcription factors upstream of the miR399-*PHO2* regulatory module in several plant species, including maize, rice, and wheat (Zhou et al., 2008, Wang et al., 2013b, Wang et al., 2013a). In maize, overexpression of *ZmPHR1* significantly increases the Pi content in shoots under low-Pi conditions (Wang et al., 2013b). By investigating the change in expression of *ZmPHR1* and its closest ortholog, *ZmMYB78*, we found that both genes showed similar transcript levels in immature leaves and mature leaves, between the *id1-crispr* mutant and wild-type plants **(Supplemental Figure S23),** suggesting that *ZmPHR1* and *ZmMYB78* are not genetically downstream of *ID1*. *PILNCR*, a Pi-deficiency-induced long non-coding RNA previously identified in maize, regulates the function of miR399 by competitively binding and sequestering the miRNA (Du et al., 2018). We also assessed the expression of *PILNCR* by qRT-PCR, and found similar expression levels in both the *id1-crispr* mutant and wild type **(Supplemental Figure S23)**. Combined with the experimental evidence showing the binding of ID1 to *ZmMIR399* promoters, these expression results support the notion of ID1 as a direct upstream regulator of *ZmMIR399* transcription. It is possible that ID1 regulates the miR399-*ZmPHO2* module independently of *ZmPHR1*.

### ID1 controls Pi homeostasis, likely through the systemic movement of miR399, in maize and possibly other crops

Plants absorb mineral nutrients from the rhizosphere via roots and the nutrients are subsequently distributed to aboveground shoots. The demand of shoots for nutrients and their supply from roots need to be properly coordinated to maintain the normal growth and development of plants. Long-distance signaling pathways can be triggered by internal or external factors as a component of adaptive responses when the nutrient status in shoots fluctuates (Forde, 2002, Atkins & Smith, 2007, Schachtman & Shin, 2007). Thus, when a change in internal nutrient status in shoots is sensed, this information is transported by long-distance signals to roots, which ensures the integration of nutritional demands in shoots with physiological changes in roots. Many nutrient uptake systems involved in nutrient assimilation and mobilization in roots are regulated by demand from the shoot via shoot-derived signals (Marschner, 1995).

In Arabidopsis, miR399 generated in shoots serves as a long-distance signal that represses *PHO2* in roots under Pi-starvation conditions, resulting in activation of Pi uptake and translocation (Lin et al., 2008, Pant et al., 2008). In maize, *ZmPHO2* transcripts in roots can be suppressed by miR399, which is expressed in the vascular systems, ultimately leading to Pi over-accumulation in shoots (Du et al., 2018). In the present study, as expected, we also found that more Pi accumulated in shoots of miR399-overexpressing transgenic maize than in the wild type (**Figure 1B**), in line with observations in Arabidopsis (Lin et al., 2008). However, in our miR399-OE transgenic maize, it is not clear whether miR399 is transported to roots to enhance Pi absorption there, because of the constitutive expression of mature miR399. In Arabidopsis, tobacco and soybean, micrografting experiments have been used to determine the shoot-to-root movement of mature miRNAs (Forde, 2002, Pant et al., 2008, Li et al., 2021). However, it is still very challenging to perform micrografting in maize, although a recent breakthrough was reported showing that the embryonic hypocotyl allows grafting in most monocotyledonous orders (Reeves et al., 2021). Accordingly, it might soon be possible to investigate the shoot-to-root movement of miR399 in maize as well.

The spatial patterns of *ID1* expression led us to speculate that its role in regulating Pi homeostasis involves the mobile miR399. The *ID1* gene is expressed exclusively in developing leaves (Colasanti et al., 1998), where *id1* is expected to repress the expression of *ZmMIR399* genes. In *id1* mutants, the repressive effect of ID1 on *ZmMIR399* transcription is released and miR399 is expected to be produced more in developing leaves. It is then likely that miR399 travels from leaves to roots, where the target gene *ZmPHO2* is downregulated and Pi absorption is enhanced to support the vigorous vegetative growth and extremely delayed flowering of the *id1* mutant. Interestingly, a mutant of the rice ortholog of the maize *ID1* gene, *early heading date2 (ehd2)*, also shows extremely delayed flowering with a giant plant architecture (Wu et al., 2008, Matsubara et al., 2008), which needs nutrients to support this extra vegetative growth. Given the conservation of the *ID1*, miR399 and *PHO2* genes (Bari et al., 2006, Chiou et al., 2006, Matsubara et al., 2008, Wu et al., 2008), this regulatory system might be conserved between maize and rice. However, unlike in rice, no *ID1* orthologs have been found in Arabidopsis (Colasanti et al., 2006). Thus, ID1 might be a monocotyledon-specific factor that orchestrates plant development and Pi acquisition to maintain growth homeostasis.

### The nutrient absorption system in maize may have been dramatically altered during domestication and expansion

Maize is well known to require large amounts of N, P and K macronutrients for optimal yield, which puts a high demand on the nutrient absorption system. After diverging from Balsas teosinte 9,000 years ago through a process of domestication and artificial selection, cultivated maize has been widely adopted as a crop plant in the fertile, highly productive plains of the temperate regions, where it is often grown with the application of fertilizer (Hastorf, 2009, Piperno et al., 2009, Sluyter & Dominguez, 2006). Compared with modern maize, teosinte grows in the wild in a variable and challenging environment in subtropical southwestern Mexico, where the plants are only subject to occasional and inadequate fertilization from recessional flooding and mineralization (Hastorf, 2009, Piperno et al., 2007, Zhu et al., 2010, Iltis et al., 1979). Thus, in addition to the dramatic morphological changes in plant size and architecture, domestication probably also caused alterations in maize metabolism and in its responses to the environment. It is also well known that most breeders involved in maize improvement preferentially focused on yield and disease resistance, which was essential to meet the demand for food due to continued growth of the human population. To increase crop yields, farmers have typically applied large quantities of synthetic fertilizer to the soil, which has often had a deleterious effect on the environment. Inevitably, the nutrient uptake system of maize was probably weakened during adaptation to such agricultural practices over long periods. Indeed, a previous report has shown that the maize progenitor, teosinte, uses nitrogen more efficiently than modern cultivated maize (Gaudin et al., 2011). Together with the evidence we present here of the weakened Pi absorption system in cultivated maize, these observations are consistent with a deterioration in nutrient absorption in modern commercial maize. In rice, *DENSE AND ERECT PANICLES 1* (*DEP1*), a G protein gene that regulates nitrogen signaling and modulation was previously reported to have been subjected to artificial selection during *Oryza sativa* spp. *japonica* domestication (Sun et al., 2014). The *DEP1* alleles from *Oryza rufipogon*, the wild ancestor of cultivated rice, confer robust resistance to nitrogen deficiency, such that the wild rice shows less sensitivity than *japonica* cultivars (Sun et al., 2014). These data in rice also point to a weakening of the nutrient absorption system in domesticated crop species. New solutions are urgently needed to simultaneously increase yields while maintaining, or preferably decreasing, fertilization to maximize the efficiency of nutrient use in crops. Otherwise, under conditions of fertilizer shortage, crop yields could be severely impacted in the future. Restoring favorable alleles from the wild ancestor, teosinte, might provide a strategy for improving the nutrient absorption system in cultivated maize and further maintaining environmentally sustainable increases in grain yield by avoiding the over-application of fertilizer.

## Materials and Methods

### Plant materials and growth conditions

Transgenic miR399-OE and miR399-STTM lines were grown in experimental fields from 2018 to 2021 in summer in Tieling (Liaoning, 41.8° N, 123.4° E), China and in winter in Sanya (Hainan, 18.4°N, 109.2°E), China, with daily management for screening of homozygous plants and bulking up of seeds. Homozygous miR399-OE, miR399-STTM and the corresponding wild-type plants were grown in the field for phenotype investigation and in a greenhouse (16 h/8 h light/dark, 25℃) to allow tissue collection for experiments.

The *id1-crispr* heterozygous mutant, created by CRISPR/Cas9, was grown in the experimental field in Tieling (Liaoning, 41.8° N, 123.4° E) or Sanya (Hainan, 18.4°N, 109.2°E) for positive mutant identification and generation of seed stocks, and in a greenhouse for genotyping and tissue collection.

### Plasmid construction and genetic transformation

For the construction of the miR399-OE binary vector, two stem-loop structures harboring different mature miR399 sequences (i.e., miR399a/c/h and miR399e/i/j) were first inserted into a primary miRNA backbone under control of the maize ubiquitin (*UBI*) promoter in the *pOT2-poly-cis-UN* cloning vector by PCR amplification. After verification by sequencing and restriction-enzyme digestion, the UBI-miR399-NOS fragment from the *pOT2-poly-cis-UN-miR399OE* construct was cloned between the *Pac*I and *Mlu*I restriction sites in the destination vector *pZZ00026-PM*, which contains the CMV35S-driven *Streptomyces hygroscopicus* phosphinothricin acetyltransferase (*BAR*) gene as a selectable marker, by a T4 ligase-mediated reaction.

The miR399-STTM binary vector was prepared following previously described procedures (Tang & Tang, 2013, Peng et al., 2018) with some modifications. In brief, the miR399-STTM fragment was first introduced into the *pOT2-poly-cis-UN* cloning vector under control of the *UBI* promoter by PCR amplification, *Swa*I digestion and T4 ligase-mediated ligation. After verification by sequencing and restriction-enzyme digestion, the UBI-miR399STTM-NOS fragment from *pOT2-poly-cis-UN-miR399STTM* was cloned between the *Pac*I and *Mlu*I sites in the binary vector *pZZ00026-PM* using T4 ligase.

To prepare the CRISPR/Cas9 knockout vector for *ID1* or *ZmPHO2*, the two target sites located in the coding region near the start codon of each gene were selected with the aid of the online tool CRISPR-P (http://crispr.hzau.edu.cn/CRISPR2/) and verified by manual alignment to the maize ‘B73’ reference genome. After restriction-enzyme digestion, the two guide RNAs (gRNAs) corresponding to the target sites were integrated into the *Btg*Z1 site under the control of the maize *U6.1* promoter and the *Bsa*I site under the control of the maize *U6.2* promoter in the *pENTR4-gRNA1* cloning vector, respectively **(Figure 3; Supplemental Figure S12)**. After verification by sequencing, the *U6-*driven gRNA cascade was cloned into the destination vector pZmCas9 by LR recombination.

After verification by sequencing, the resulting binary construct containing either miR399-OE or miR399-STTM and CRISPR/Cas9 was introduced into *Agrobacterium tumefaciens* EHA105 for genetic transformation. To identify mutation events, genomic fragments covering both target sites were amplified from the T0 CRISPR/Cas9 plants by PCR. After gel purification, the resulting PCR products were cloned into the pEASY-Blunt cloning vector (TransGen, Beijing, China) and at least ten clones per PCR product were analyzed by Sanger sequencing. Finally, one and three independent T0 lines with large fragment deletions at the target sites were identified for *ID1* and *ZmPHO2*, respectively (**Figure 3; Supplemental Figure S12**).

### Quantification of total phosphorus content

Shoots and roots of transgenic miR399-OE and miR399-STTM plants were collected separately with three biological replicates at the seedling stage. For each replicate, tissues from more than five plants were first pooled and incubated at 105°C for 30 min. The samples were then dried at 65°C for 3 d, weighed and milled to a fine powder. Portions (∼100 µg) were then digested in 5 mL H_2_SO_4_-H_2_O_2_ at 300°C until the solution became clear. The total phosphorus content was determined by the vanadomolybdophosphoric acid colorimetric method. The same protocol was also used to determine the total phosphorus content in mature leaf, immature leaf, root, stem and leaf sheath in the *id1-crispr* mutant and wild-type plants.

### Measurement of relative chlorophyll content

At 15 days after pollination, when the premature aging syndrome appeared, the relative chlorophyll contents of ear leaves were measured with the mobile device, SPAD 502 Plus Chlorophyll Meter (Speactrum Technologies, Inc.), which is widely used for the rapid, accurate and non-destructive measurement of leaf chlorophyll concentrations. Measurements with the SPAD 502 Plus meter produce relative SPAD (soil plant analysis development) values that are proportional to the amount of chlorophyll present. Three different regions of the leaf, base, middle and tip, were chosen to measure relative chlorophyll content. More than 30 independent plants were used for analysis within an experiment.

### Gene expression analysis with qRT-PCR

Total RNA was extracted from the mature or immature leaves of *id1-crispr* mutant, miR399-OE transgenic plants, miR399-STTM transgenic plants and their corresponding wild-type controls using TRIzol reagent (Invitrogen, USA). Genomic DNA removal and first-strand cDNA synthesis were carried out using a TAKARA first-strand cDNA synthesis kit (TaKaRa, Japan). Quantitative real-time PCR (qRT-PCR) was performed using SYBR Green detection and relative gene expression was determined with the 2^-△△Ct^ method (Livak and Schmittgen, 2001) from three biological replicates. *ZmUbiquitin* was used as the internal control for quantitation of mRNA levels. Primers used for qRT-PCR are listed in **Supplemental Dataset S4**.

### Small RNA northern blot

Total RNA was isolated from mature and immature leaves of the *id1-crispr* mutant and wild type. Three biological replicates were collected for all samples with at least five leaves pooled for each genotype. Total RNA was used for northern blot analysis to assess the accumulation of mature miR399 in the *id1-crispr* mutant. In brief, a 15% urea PAGE gel was made from urea gel concentrate, urea diluent and urea gel buffer (National Diagnostics). Ten percent ammonium persulfate (APS; 100 µl) and 10 µl TEMED were added per 15 ml gel mixture to polymerize the gel. A minimum of 10 µg RNA per sample was loaded into a 15% urea PAGE gel and resolved at 120 V for 1 h. RNA was transferred from the gel to a nitrocellulose membrane (Bio-Rad) using a semi-dry transfer apparatus (Bio-Rad). Specific northern blot probes were labeled with 5’-biotin and hybridized with the membrane. A probe complementary to U6 was used as an internal control. Hybridization was performed for 16 h at 55°C followed by washes. Signals were detected using the Chemiluminescent Nucleic Acid Detection Module (Thermo Fisher, 89880) with a chemiluminescence imaging system (Clinx Science Instruments Co. Ltd., China). Probe sequences can be found in **Supplemental Dataset S4.**

### Recombinant protein expression and purification, electrophoretic mobility shift assay (EMSA) and *in vitro* DNA pull-down assay

To construct the MBP-ID1 expression vector, the coding sequence (CDS) of *ID1* was first amplified from cDNA using primers ID1-MBP-F and ID1-MBP-R **(Supplemental Dataset S4)**. The PCR product was then cloned into a version of the *pMSCG7* vector with the maltose binding protein (MBP) coding sequence at the N-terminal end of the cloning site using Gibson assembly; the resulting construct was validated by Sanger sequencing. The empty expression vector and the version containing *ID1* CDS were introduced separately into RosettaTM DE3 *E. coli* (Sigma-Aldrich, 70954) for protein expression and purification. Transformed *E. coli* clones were first induced with 0.1 mM IPTG (Thermo Fisher Scientific, 15529019) at 16°C for 16 h. The bacteria pellets were collected at 4°C by centrifugation at 5,000 rpm and re-suspended in lysis buffer [100 mM Tris-HCl, pH7.5; 500 mM NaCl; 5 mM 2-mercaptoethanol; 5% glycerol; 20 mM imidazole; 0.1% CA630; 1 mM PMSF; 1 x EDTA-free protease inhibitor cocktail]. After sonication, the recombinant protein was purified with Profinity™ IMAC Resin, Ni-charged (Bio-Rad, 1560133). The eluted protein concentration was estimated using a Bradford assay (Quick Start™ Bradford 1x Dye Reagent #5000205; Bio-Rad).

For EMSA, a 20 µl reaction mixture containing 10 x binding buffer [100 mM Tris-HCl, pH 7.5; 500 mM KCl; 10 mM DTT], 20-fmol biotin-labeled probe **(Supplemental Dataset S4)**, recombinant proteins (∼ 1 µg), and varying amounts of unlabeled probes as competitors was incubated at room temperature for 1 h. DNA-protein complexes were resolved in a 6% polyacrylamide gel and transferred to a charged Hybond-N^+^ membrane (GE Healthcare). The membrane was cross-linked by UV (1200 mJ), probed with stabilized streptavidin-horseradish peroxidase (HRP) and subsequently detected using the Chemiluminescent Nucleic Acid Detection Module Kit (Thermo Fisher). Membranes were imaged using a ChemiDoc XRS+ camera (BioRad).

For the *in vitro* DNA pull-down assay, ∼10 pmol of the biotin-labeled probes used in the EMSA experiments mentioned above were first incubated with 10 µl DynabeadsTM M-280 streptavidin magnetic beads (Thermo Fisher Scientific) for 15 min at room temperature in 1 x B&W buffer [5 mM Tris-HCl (pH 7.4), 0.5 mM EDTA, 1 M NaCl]. Probe-bound beads were then incubated with the same amount of MBP or MBP-ID1 protein (∼ 100 ng) in 100 µl binding buffer [10 mM Tris·HCl (pH 7.5), 50 mM KCl, 5 mM MgCl_2_, 2.5% glycerol, 0.05% CA630] supplemented with 10 µg BSA for 1 h at 4°C. The precipitates and one-fifth of the supernatant were boiled in SDS loading buffer and subjected to western blot analysis using an anti-MBP monoclonal antibody (New England Biolabs, E8032S, 1:200 dilution), followed by anti-mouse IgG-HRP (Sigma, A4416, 1:5,000 dilution).

### RNA-seq library preparation, sequencing and data processing

The transgenic lines, including miR399-OE#2 and miR399-STTM#2, and the corresponding wild type, were used for RNA sequencing with three biological replicates. Mature leaves were collected at the floral transition stage and at least five leaves were pooled for each replicate. Total RNA extraction was performed using the Trizol reagent and genomic DNA was removed. Total RNA quality was verified using an Agilent 2100 Bioanalyzer (Agilent Tehnologies, Santa Clara, USA) before library preparation. Sequencing libraries were prepared with a NEBNext Ultra II RNA Library Prep Kit for Illumina (NEB) and sequenced on an Illumina HiSeqX10 platform using a 150-bp paired-end sequencing strategy at Novogene Co., Ltd. (Beijing, China). Approximately 25 million high-quality 150-bp paired-end reads were generated from each library.

The raw data were first examined for library quality using the FastQC program (https://github.com/s-andrews/FastQC). Reads that passed the quality check were analyzed using the ‘pRNASeqTools mrna’ pipeline (https://github.com/grubbybio/pRNASeqTools/), which integrates the trimming of low-quality nucleotides, reads alignment, reads/fragments counting and differential gene expression analysis. In brief, trimmed reads of high quality were mapped to the B73 genome (https://www.maizegdb.org/) using STAR 2.7.3a with the parameters “--alignIntronMax 5000 –outSAMmultNmax 1 --outFilterMultimapNmax 50 --outFilterMismatchNoverLmax 0.1” (Dobin et al., 2013). Then, featureCounts (v2.0.0) was employed to count reads mapped to exon regions for each gene with the parameters “-p -B –C -O -s 0” (Liao et al., 2014). Differentially expressed genes (DEGs) were identified with fold change (RPM) ≥ 2 and FDR < 0.05 using DESeq2 (Love et al., 2014). Gene Ontology and KEGG enrichment analysis of the DEGs was performed using g:Profiler, a free online platform (https://biit.cs.ut.ee/gprofiler/gost) (Raudvere et al., 2019). GO terms with FDR ≤ 0.05 were retained and deemed significantly enriched.

### Small RNA-seq data analysis

The raw small RNA-seq data for the *id1-m1* mutant (accession number PRJNA439244) were downloaded from the National Center for Biotechnology Information Sequence Read Archive (Minow et al., 2018). The 3′ adaptor sequence was first detected and trimmed, and then size selection (from 18-nt to 42-nt) was carried out for adaptor-trimmed reads using Cutadapt v1.15 (Martin, 2011). The retained reads were mapped to the AGPv4 genome for B73 maize (https://www.maizegdb.org/) using ShortStack v3.8.5 (Johnson et al., 2016). Information on annotated maize miRNAs was adopted from both miRBase v21 (http://www.mirbase.org/) and miRNEST 2.0 (http://rhesus.amu.edu.pl/mirnest/copy/). Adaptor-trimmed reads that mapped to specific miRNAs were counted and summarized separately for each size class ranging from 18-nt to 27-nt. Normalization was performed by calculating the reads-per-million (RPM) value for each size class, and comparison was carried out using the R package DESeq2 (Love et al., 2014).

### Dual-luciferase transient expression assay in maize protoplasts

To test the repression of *MIR399c* and *MIR399j* expression by ID1 *in vivo*, we performed a dual-luciferase transient expression assay in maize protoplasts. The 2-kb promoter fragments of *MIR399c* and *MIR399j* were cloned into the *pGreen II 0800-LUC* vector, which carries a minimal promoter from the cauliflower mosaic virus (mpCaMV) upstream of the luciferase (*LUC*) coding sequence to drive *LUC* reporter gene transcription (**Figure 5B**). The *Renilla* luciferase gene (*REN*) under the control of the 35S promoter in the *pGreen II 0800-LUC* vector was used as an internal control to correct for protoplast transfection efficiency differences. The *ID1* coding sequence was cloned into the *pGreen II 62-SK* vector as effector (**Figure 5B**).

Mesophyll protoplasts were isolated from leaves of 12-day-old etiolated B73 seedlings following the method described previously (Yoo et al., 2007). The *ID1-62-SK* and *pMIR399c-0800-LUC* or *pMIR399j-0800-LUC* plasmids were co-transformed into the mesophyll protoplasts using the previously described polyethylene glycol method (Yoo et al., 2007, Hellens et al., 2005). After incubation in the dark for 12-16 h, LUC and REN activities were measured using the Dual-Luciferase Reporter Assay System (Promega) according to the manufacturer’s instructions. Relative firefly LUC activity was calculated by normalizing LUC activity against REN activity. Six technical replicates were performed per construct combination - the same batch of protoplasts was split into six different tubes as the technical replicates, which were transformed with a combination of plasmids. All experiments were independently repeated three times. Primers used for plasmid construction are shown in **Supplemental Dataset S4**.

### Agrobacterium-mediated infiltration and fluorescence microscopy in *Nicotiana benthamiana* leaves

To construct *35S::ID1-EYFP*, the *ID1* coding region was PCR-amplified from genomic DNA using primers ID1-TSK-F and ID1-TSK-R **(Supplemental Dataset 4)**. The PCR product was then cloned into *TSK108*, a pENTRY-D-topo-based Gateway entry vector. The sequence of the pENTR clone was validated by DNA sequencing and recombined with the binary vector pGWB641, which contains an EYFP tag at its C-terminal end, by Gateway cloning. Similarly, the vector *35S::FIB2-RFP* was prepared with *pGWB654* as a nucleolus marker.

For subcellular localization in *Nicotiana benthamiana* leaf epidermal cells, the *Agrobacterium tumefaciens* strain GV3101 containing the plasmid constructs, *35S::ID1-EYFP* and *35S::FIB2-RFP*, was grown overnight at 28℃ with constant shaking at 200 rpm. The *Agrobacterium* cells were collected by centrifugation at 5,000 rpm. The supernatant was discarded and the pellet was resuspended in infiltration buffer [10 mM MES (pH 5.6), 10 mM MgCl_2_, 150 mM acetosyringone] at OD=1.0. Subsequently, the *Agrobacterium* resuspensions of *35S::ID1-EYFP* and *35S::FIB2-RFP* were mixed evenly in equal volumes and co-infiltrated into four-week-old leaves of *N. benthamiana*. After infiltration, plants were placed under long-day conditions (16 h light/8 h dark, 22℃) for ∼48 h, and EYFP and RFP fluorescence was observed via confocal microscopy using a Zeiss 710 laser scanning microscope (Carl Zeiss, Oberkochen, Germany). To stain the nucleus, the fluorescent stain, 4’, 6-diamidino-2-phenylindole (DAPI), was infiltrated into the leaves, and DAPI fluorescence was observed at 20 min after infiltration via confocal microscopy.

### Nucleotide diversity analysis and test for selection

The third generation haplotype map data of *Zea mays* (HapMap 3: https://www.panzea.org/) (Bukowski et al., 2018) was downloaded and used to calculate nucleotide diversity along the 10 kb upstream and downstream regions of *MIR399* family genes and *ZmPHO2* in maize, landraces and teosinte. The population genomic analysis was done with the R package PopGenome (Pfeifer et al., 2014). Sliding windows (window size = 1,000 bp, step size = 100 bp) across the target regions were generated using the function ‘sliding.window.transform’. Nucleotide diversity (*π*) in each sliding window was calculated for three groups, including tropical maize, temperate maize and teosinte, using the function ‘diversity.stats’. For the analysis of the re-sequenced ∼1.7-kb region in *ZmPHO2*, multiple sequence alignments were performed using BioEdit v.7.1.3.0 with some manual editions when necessary. Nucleotide diversity was calculated using DnaSP v.5.10.00 with a sliding window size of 100-bp. Coalescent simulations were employed to evaluate whether the loss of nucleotide diversity observed in maize relative to that in teosinte could be explained by a domestication bottleneck alone using the Hudson’s ms program (Hudson, 2002). All parameters in the model were set according to previously established values (Xu et al., 2017, Tian et al., 2009, Wright et al., 2005, Liang et al., 2019). The population mutation and population recombination parameters were estimated from the teosinte sequences. A total of 10,000 coalescent simulations were performed.

## Data availability

The raw RNA-seq data in this study have been submitted to the GenBank database under accession number PRJNA819017.

## Acknowledgements

This research was supported by the National Natural Science Foundation of China (31900446), Guangdong Innovation Research Team Foundation (2014ZT05S078), China Postdoctoral Science Foundation (2018M640823), and Shenzhen Basic Research General Project (JCYJ20190808112207542).

## Conflict of interest statement

The authors declare no conflict of interest.

## Author contributions

X.W., X.C. and L.L. designed and supervised the study; X.W., D.Y., YC.L. (Yanchun Liu), YM.L. (Yameng Liang), J.H., X.Y., R.H., and Q.X. conducted laboratory experiments, field work and data analysis; YM.L., H. J., and F.T. created the *id1-crispr* mutant by CRISPR/Cas9 technology and near-isogenic lines, and provided the DIBOA and DIMBOA standards; X.W. performed the RNA-seq and small RNA-seq data analysis and interpretation; X.W., X.C. and L.L. wrote the manuscript. All authors read and revised the manuscript.

## References

Aguirre-Liguori JA, Gaut BS, Jaramillo-Correa JP, et al., 2019. Divergence with gene flow is driven by local adaptation to temperature and soil phosphorus concentration in teosinte subspecies *(Zea mays parviglumis* and *Zea mays mexicana*). Mol Ecol 28, 2814–30.

Andorf C, Beavis WD, Hufford M, et al., 2019. Technological advances in maize breeding: past, present and future. Theor Appl Genet 132, 817–49.

Atkins CA, Smith PMC, 2007. Translocation in legumes: assimilates, nutrients, and signaling molecules. Plant Physiol 144, 550–61.

Aung K, Lin S-I, Wu C-C, Huang Y-T, Su C-L, Chiou T-J, 2006. *pho2*, a phosphate overaccumulator, is caused by a nonsense mutation in a microRNA399 target gene. Plant Physiol 141, 1000–11.

Baker A, Ceasar SA, Palmer AJ, et al., 2015. Replace, reuse, recycle: improving the sustainable use of phosphorus by plants. J Exp Bot 66, 3523–40.

Bari R, Datt Pant B, Stitt M, Scheible W-R, 2006. *PHO2*, microRNA399, and *PHR1* define a phosphate-signaling pathway in plants. Plant Physiol 141, 988–99.

Belhaj K, Chaparro-Garcia A, Kamoun S, Patron NJ, Nekrasov V, 2015. Editing plant genomes with CRISPR/Cas9. Curr Opin Biotechnol 32, 76–84.

Bukowski R, Guo X, Lu Y, et al., 2018. Construction of the third-generation *Zea mays* haplotype map. Gigascience 7, gix134.

Bustos R, Castrillo G, Linhares F, et al., 2010. A central regulatory system largely controls transcriptional activation and repression responses to phosphate starvation in Arabidopsis. PLoS Genet 6, e1001102.

Campos-Soriano L, Bundó M, Bach-Pages M, Chiang SF, Chiou TJ, San Segundo B, 2020. Phosphate excess increases susceptibility to pathogen infection in rice. Mol Plant Pathol 21, 555–70.

Chen Z-H, Nimmo GA, Jenkins GI, Nimmo HG, 2007. *BHLH32* modulates several biochemical and morphological processes that respond to Pi starvation in Arabidopsis. Biochem J 405, 191–8.

Chiou T-J, Aung K, Lin S-I, Wu C-C, Chiang S-F, Su C-L, 2006. Regulation of phosphate homeostasis by microRNA in Arabidopsis. Plant Cell 18, 412–21.

Colasanti J, Sundaresan V, 2000. ‘Florigen’ enters the molecular age: long-distance signals that cause plants to flower. Trends Biochem Sci 25, 236–40.

Colasanti J, Tremblay R, Wong AY, Coneva V, Kozaki A, Mable BK, 2006. The maize INDETERMINATE1 flowering time regulator defines a highly conserved zinc finger protein family in higher plants. BMC Genom 7, 1–17.

Colasanti J, Yuan Z, Sundaresan V, 1998. The indeterminate gene encodes a zinc finger protein and regulates a leaf-generated signal required for the transition to flowering in maize. Cell 93, 593–603.

Coneva V, Zhu T, Colasanti J, 2007. Expression differences between normal and *indeterminate1* maize suggest downstream targets of ID1, a floral transition regulator in maize. J Exp Bot 58, 3679–93.

Cordell D, Rosemarin A, Schröder JJ, Smit A, 2011. Towards global phosphorus security: A systems framework for phosphorus recovery and reuse options. Chemosphere 84, 747–58.

Delhaize E, Randall PJ, 1995. Characterization of a phosphate-accumulator mutant of *Arabidopsis thaliana*. Plant Physiol 107, 207–13.

Devaiah BN, Karthikeyan AS, Raghothama KG, 2007a. WRKY75 transcription factor is a modulator of phosphate acquisition and root development in Arabidopsis. Plant Physiol 143, 1789–801.

Devaiah BN, Madhuvanthi R, Karthikeyan AS, Raghothama KG, 2009. Phosphate starvation responses and gibberellic acid biosynthesis are regulated by the MYB62 transcription factor in Arabidopsis. Mol Plant 2, 43–58.

Devaiah BN, Nagarajan VK, Raghothama KG, 2007b. Phosphate homeostasis and root development in Arabidopsis are synchronized by the zinc finger transcription factor *ZAT6*. Plant Physiol 145, 147–59.

Devaiah BN, Raghothama KG, 2007. Transcriptional regulation of Pi starvation responses by WRKY75. Plant Signal Behav 2, 424–5.

Dobin A, Davis CA, Schlesinger F, et al., 2013. STAR: ultrafast universal RNA-seq aligner. Bioinformatics 29, 15–21.

Doudna JA, Charpentier E, 2014. The new frontier of genome engineering with CRISPR-Cas9. Science 346,1258096

Du Q, Wang K, Zou C, Xu C, Li W-X, 2018. The PILNCR1-miR399 regulatory module is important for low phosphate tolerance in maize. Plant Physiol 177, 1743–53.

Eyre-Walker A, Gaut RL, Hilton H, Feldman DL, Gaut BS, 1998. Investigation of the bottleneck leading to the domestication of maize. Proc Natl Acad Sci U S A 95, 4441–6.

Fang C, Luo J, 2019. Metabolic GWAS-based dissection of genetic bases underlying the diversity of plant metabolism. The Plant J 97, 91–100.

Forde BG, 2002. The role of long-distance signalling in plant responses to nitrate and other nutrients. J Exp Bot 53, 39–43.

Franco-Zorrilla JM, Valli A, Todesco M, et al., 2007. Target mimicry provides a new mechanism for regulation of microRNA activity. Nat Genet 39, 1033–7.

Frey M, Chomet P, Glawischnig E, et al., 1997. Analysis of a chemical plant defense mechanism in grasses. Science 277, 696–9.

Frey M, Schullehner K, Dick R, Fiesselmann A, Gierl A, 2009. Benzoxazinoid biosynthesis, a model for evolution of secondary metabolic pathways in plants. Phytochemistry 70, 1645–51.

Fujii H, Chiou T-J, Lin S-I, Aung K, Zhu J-K, 2005. A miRNA involved in phosphate-starvation response in Arabidopsis. Curr Bio 15, 2038–43.

Gaudin AC, Mcclymont SA, Raizada MN, 2011. The nitrogen adaptation strategy of the wild teosinte ancestor of modern maize, Zea mays subsp. parviglumis. Crop Sci 51, 2780–95.

Godfrey O, 2010. Improved inputs use and productivity in Uganda’s maize sub-sector. No. 677-2016-46694

Goedert W, 1983. Management of the Cerrado soils of Brazil: a review. J Soil Sci 34, 405–28.

Hackenberg M, Shi B-J, Gustafson P, Langridge P, 2013. Characterization of phosphorus-regulated miR399 and miR827 and their isomirs in barley under phosphorus-sufficient and phosphorus-deficient conditions. BMC Plant Bio 13, 1–17.

Hastorf CA, 2009. Rio Balsas most likely region for maize domestication. Proc Natl Acad Sci U S A 106, 4957–8.

Hellens RP, Allan AC, Friel EN, et al., 2005. Transient expression vectors for functional genomics, quantification of promoter activity and RNA silencing in plants. Plant methods 1, 13.

Hu B, Zhu C, Li F, et al., 2011. *LEAF TIP NECROSIS1* plays a pivotal role in the regulation of multiple phosphate starvation responses in rice. Plant Physiol 156, 1101–15.

Hudson RR, 2002. Generating samples under a Wright-Fisher neutral model of genetic variation. Bioinformatics 18, 337–8.

Iltis HH, Doebley JF, Guzmán R, Pazy B, 1979. *Zea diploperennis* (Gramineae): a new teosinte from Mexico. Science 203, 186–8.

Johnson NR, Yeoh JM, Coruh C, Axtell MJ, 2016. Improved placement of multi-mapping small RNAs. G3: Genes, Genomes, Genet 6, 2103–11.

Kirkby EA, Johnston AEJ, 2008. Soil and fertilizer phosphorus in relation to crop nutrition. The ecophysiology of plant-phosphorus interactions, 177–223.

Kozaki A, Hake S, Colasanti J, 2004. The maize ID1 flowering time regulator is a zinc finger protein with novel DNA binding properties. Nucleic Acids Res 32, 1710–20.

Kremling KA, Chen S-Y, Su M-H, et al., 2018. Dysregulation of expression correlates with rare-allele burden and fitness loss in maize. Nature 555, 520–3.

López-Arredondo DL, Leyva-González MA, González-Morales SI, López-Bucio J, Herrera-Estrella L, 2014. Phosphate nutrition: improving low-phosphate tolerance in crops. Annu Rev Plant Biol 65, 95–123.

Li S, Wang X, Xu W, et al., 2021. Unidirectional movement of small RNAs from shoots to roots in interspecific heterografts. Nature Plants 7, 50–9.

Li S, Ying Y, Secco D, et al., 2017. Molecular interaction between PHO2 and GIGANTEA reveals a new crosstalk between flowering time and phosphate homeostasis in *Oryza sativa*. Plant Cell Environ 40, 1487–99.

Liang Y, Liu Q, Wang X, et al., 2019. *ZmMADS69* functions as a flowering activator through the *ZmRap2.7-ZCN 8* regulatory module and contributes to maize flowering time adaptation. New Phytol 221, 2335–47.

Liao Y, Smyth GK, Shi W, 2014. featureCounts: an efficient general purpose program for assigning sequence reads to genomic features. Bioinformatics 30, 923–30.

Lin S-I, Chiang S-F, Lin W-Y, et al., 2008. Regulatory network of microRNA399 and *PHO2* by systemic signaling. Plant Physiol 147, 732–46.

Liu T-Y, Huang T-K, Tseng C-Y, et al., 2012. *PHO2*-dependent degradation of PHO1 modulates phosphate homeostasis in Arabidopsis. Plant Cell 24, 2168–83.

Love MI, Huber W, Anders S, 2014. Moderated estimation of fold change and dispersion for RNA-seq data with DESeq2. Genome Biol 15, 1–21.

Luo B, Ma P, Nie Z, et al., 2019. Metabolite profiling and genome-wide association studies reveal response mechanisms of phosphorus deficiency in maize seedling. Plant J 97, 947–69.

Lynch JP, Brown KM, 2001. Topsoil foraging-an architectural adaptation of plants to low phosphorus availability. Plant Soil 237, 225–37.

Ma X, Zhang Q, Zhu Q, et al., 2015. A robust CRISPR/Cas9 system for convenient, high-efficiency multiplex genome editing in monocot and dicot plants. Mol Plant 8, 1274–84.

Maeda Y, Konishi M, Kiba T, et al., 2018. A NIGT1-centred transcriptional cascade regulates nitrate signalling and incorporates phosphorus starvation signals in Arabidopsis. Nat Commun 9, 1–14.

Marschner H, 1995. Mineral nutrition of higher plants. Academic Press. Inc*., San Diago*.

Martín-Robles N, Morente-López J, Freschet GT, Poorter H, Roumet C, Milla R, 2019. Root traits of herbaceous crops: Pre-adaptation to cultivation or evolution under domestication? Funct Ecol 33, 273–85.

Martin M, 2011. Cutadapt removes adapter sequences from high-throughput sequencing reads. EMBnet J 17, 10–2.

Matsubara K, Yamanouchi U, Wang ZX, Minobe Y, Izawa T, Yano M, 2008. *Ehd2*, a rice ortholog of the maize *INDETERMINATE1* gene, promotes flowering by up-regulating *Ehd1*. Plant Physiol 148, 1425–35.

Matsuoka Y, Vigouroux Y, Goodman MM, Sanchez J, Buckler E, Doebley J, 2002. A single domestication for maize shown by multilocus microsatellite genotyping. Proc Natl Acad Sci U S A 99, 6080–4.

Minow MaA, vila LM, Turner K, et al., 2018. Distinct gene networks modulate floral induction of autonomous maize and photoperiod-dependent teosinte. J Exp Bot 69, 2937–52.

Pant BD, Buhtz A, Kehr J, Scheible WR, 2008. MicroRNA399 is a long-distance signal for the regulation of plant phosphate homeostasis. The Plant J 53, 731–8.

Peng T, Qiao M, Liu H, et al., 2018. A resource for inactivation of microRNAs using short tandem target mimic technology in model and crop plants. Mol Plant 11, 1400–17.

Pfeifer B, Wittelsbürger U, Ramos-Onsins SE, Lercher MJ, 2014. PopGenome: an efficient Swiss army knife for population genomic analyses in R. Mol Biol Evo 31, 1929–36.

Piperno DR, Moreno JE, Iriarte J, et al., 2007. Late Pleistocene and Holocene environmental history of the Iguala Valley, central Balsas watershed of Mexico. Proc Natl Acad Sci U S A 104, 11874–81.

Piperno DR, Ranere AJ, Holst I, Iriarte J, Dickau R, 2009. Starch grain and phytolith evidence for early ninth millennium BP maize from the Central Balsas River Valley, Mexico. Proc Natl Acad Sci U S A 106, 5019–24.

Raghothama K, 1999. Phosphate acquisition. Annu Rev Plant Bio 50, 665–93.

Raudvere U, Kolberg L, Kuzmin I, et al., 2019. g: Profiler: a web server for functional enrichment analysis and conversions of gene lists (2019 update). Nucleic Acids Res 47, W191–W8.

Reeves G, Tripathi A, Singh P, et al., 2021. Monocotyledonous plants graft at the embryonic root-shoot interface. Nature, 1-7.

Rubio V, Linhares F, Solano R, et al., 2001. A conserved MYB transcription factor involved in phosphate starvation signaling both in vascular plants and in unicellular algae. Genes Dev 15, 2122–33.

Schachtman DP, Shin R, 2007. Nutrient sensing and signaling: NPKS. Annu. Rev. Plant Biol. 58, 47–69.

Sega P, Kruszka K, Szewc Ł, Szweykowska-Kulińska Z, Pacak A, 2020. Identification of transcription factors that bind to the 5’-UTR of the barley *PHO2* gene. Plant Mol Biol 102, 73–88.

Shin H, Shin HS, Chen R, Harrison MJ, 2006. Loss of *At4* function impacts phosphate distribution between the roots and the shoots during phosphate starvation. The Plant J 45, 712–26.

Singleton WR, 1946. Inheritance of indeterminate growth in maize. Journal of Heredity 37, 61–4.

Sluyter A, Dominguez G, 2006. Early maize (*Zea mays L.*) cultivation in Mexico: Dating sedimentary pollen records and its implications. Proc Natl Acad Sci U S A 103, 1147–51.

Stefanovic A, Ribot C, Rouached H, et al., 2007. Members of the *PHO1* gene family show limited functional redundancy in phosphate transfer to the shoot, and are regulated by phosphate deficiency via distinct pathways. The Plant J 50, 982–94.

Sun H, Qian Q, Wu K, et al., 2014. Heterotrimeric G proteins regulate nitrogen-use efficiency in rice. Nat Genet 46, 652–6.

Tang G, Tang X, 2013. Short tandem target mimic: a long journey to the engineered molecular landmine for selective destruction/blockage of microRNAs in plants and animals. J Genet Genom 40, 291–6.

Tang G, Yan J, Gu Y, et al., 2012. Construction of short tandem target mimic (STTM) to block the functions of plant and animal microRNAs. Methods 58, 118–25.

Tian F, Stevens NM, Buckler ES, 2009. Tracking footprints of maize domestication and evidence for a massive selective sweep on chromosome 10. Proc Natl Acad Sci U S A 106, 9979–86.

Vance CP, Uhde-Stone C, Allan DL, 2003. Phosphorus acquisition and use: critical adaptations by plants for securing a nonrenewable resource. New Phytol 157, 423–47.

Wang C, Ying S, Huang H, Li K, Wu P, Shou H, 2009. Involvement of *OsSPX1* in phosphate homeostasis in rice. The Plant J 57, 895–904.

Wang J, Sun J, Miao J, et al., 2013a. A phosphate starvation response regulator *Ta-PHR1* is involved in phosphate signalling and increases grain yield in wheat. Ann Bot 111, 1139–53.

Wang X, Bai J, Liu H, Sun Y, Shi X, Ren Z, 2013b. Overexpression of a maize transcription factor ZmPHR1 improves shoot inorganic phosphate content and growth of Arabidopsis under low-phosphate conditions. Plant Mol Biol Rep 31, 665–77.

Wang X, Chen Q, Wu Y, et al., 2018. Genome-wide analysis of transcriptional variability in a large maize-teosinte population. Mol Plant 11, 443–59.

Wright SI, Bi IV, Schroeder SG, et al., 2005. The effects of artificial selection on the maize genome. Science 308, 1310–4.

Wu C, You C, Li C, et al., 2008. RID1, encoding a Cys2/His2-type zinc finger transcription factor, acts as a master switch from vegetative to floral development in rice. Proc Natl Acad Sci U S A 105, 12915–20.

Xu F, Liu Q, Chen L, et al., 2013. Genome-wide identification of soybean microRNAs and their targets reveals their organ-specificity and responses to phosphate starvation. BMC Genom 14, 1–30.

Xu G, Wang X, Huang C, et al., 2017. Complex genetic architecture underlies maize tassel domestication. New Phytol 214, 852–64.

Yoo S-D, Cho Y-H, Sheen J, 2007. Arabidopsis mesophyll protoplasts: a versatile cell system for transient gene expression analysis. Nat Protoc 2, 1565–72.

Zhang L, Chia J-M, Kumari S, et al., 2009. A genome-wide characterization of microRNA genes in maize. PLoS Genet 5, e1000716.

Zhou J, Jiao F, Wu Z, et al., 2008. *OsPHR2* is involved in phosphate-starvation signaling and excessive phosphate accumulation in shoots of plants. Plant Physiol 146, 1673–86.

Zhu J, Zhang C, Lynch JP, 2010. The utility of phenotypic plasticity of root hair length for phosphorus acquisition. Funct Plant Biol 37, 313–22.

